# Cephalopod Retinal Development Shows Vertebrate-like Mechanisms of Neurogenesis

**DOI:** 10.1101/2021.10.28.466353

**Authors:** Francesca Napoli, Christina M. Daly, Stephanie Neal, Kyle J. McCulloch, Alexandra Zaloga, Alicia Liu, Kristen M. Koenig

## Abstract

Neurogenesis, the regulation of cellular proliferation and differentiation in the developing nervous system, is the process that underlies the diversity of size and cell type found in animal nervous systems. Our understanding of how this process has evolved is limited because of the lack of high resolution data and live-imaging methods across species. The retina is a classic model for the study of neurogenesis in vertebrates and live-imaging of the retina has shown that during development, progenitor cells are organized in a pseudostratified neuroepithelium and nuclei migrate in coordination with the cell cycle along the apicobasal axis of the cell, a process called interkinetic nuclear migration. Eventually cells delaminate and differentiate within the boundaries of the epithelium. This process has been considered unique to vertebrates and thought to be important in maintaining organization during the development of a complex nervous system. Coleoid cephalopods, including squid, cuttlefish and octopus, have the largest nervous system of any invertebrate and convergently-evolved camera-type eyes, making them a compelling comparative system to vertebrates. Here we have pioneered live-imaging techniques to show that the squid, *Doryteuthis pealeii*, displays cellular mechanisms during cephalopod retinal neurogenesis that are hallmarks of vertebrate processes. We find that retinal progenitor cells in the squid undergo interkinetic nuclear migration until they exit the cell cycle, we identify retinal organization corresponding to progenitor, post-mitotic and differentiated cells, and we find that Notch signaling regulates this process. With cephalopods and vertebrates having diverged 550 million years ago, these results suggest that mechanisms thought to be unique to vertebrates may be common to highly proliferative neurogenic primordia contributing to a large nervous system.

## Introduction

Coleoid cephalopods (e.g. squid, cuttlefish and octopus) are charismatic invertebrates known for their expansive behavioral repertoire and large nervous system. One of the striking features of the cephalopod nervous system is their camera-type eyes, which are one of the most acute visual organs among animals (Gagnon et al., 2013; Packard, 1972). This complexity in cephalopods is independently evolved from similar features found in vertebrate species. Although vertebrates have been well studied, the developmental changes that underlie the evolution of large, complex nervous systems are not well understood. Observations in traditional model organisms show that diversity in nervous system size and functionality across animals is a product of diversity in developmental processes (Hartenstein & Stollewerk, 2015). We know that in some cases neurogenesis is highly regulated with invariant numbers of precursors with fixed lineages (i.e. *C. elegans*), and in others cell lineages are stochastic or plastic (i.e. the vertebrate retina) (Bertrand & Hobert, 2010; Zechner et al., 2020); (Hartenstein & Stollewerk, 2015). Live-imaging has changed our perspective on neurodifferentiation from a primarily genetic process to a cell biological process, defined by orchestrated cell behaviors and tissue architecture. However, *in vivo* observations of neurogenesis have been limited to vertebrates and *Drosophila*. To elucidate how changes in neurogenesis may contribute to the evolution of large and complex nervous systems, we sought to understand neurodifferentiation in the cephalopod. The visual system has proven to be a powerful context to learn fundamental aspects of neural development in vertebrates and *Drosophila* and therefore we have chosen to focus our investigations on retinal differentiation in the squid *Doryteuthis pealeii*.

The adult cephalopod retina is composed of two layers divided by the basal membrane (Figure 1A). Photoreceptor cell bodies are found posterior (further from the lens) to the basal membrane and support cells are found anterior (closer to the lens). The photoreceptor cells extend projections to the anterior of the retina to form the outer segment layer and extend axons out of the posterior retina to synapse directly on the optic lobe (Yamamoto et al., 1965; Young, 1974). During development, photoreceptor cell bodies are first identified behind the basal membrane at Arnold embryonic stage 27, and these cells are no longer proliferative (Arnold, 1965; Koenig et al., 2016). This is the earliest evidence of differentiation in the retina. The support cells, in the anterior of the retina, also extend processes into the outer segment (Yamamoto et al., 1965). Although the eye is functional at hatching, the support cells continue to incorporate BrdU (Koenig et al., 2016). The animal continues to grow post-hatching and proliferating support cells may include a long-term retinal stem cell population.

**Figure 1:**
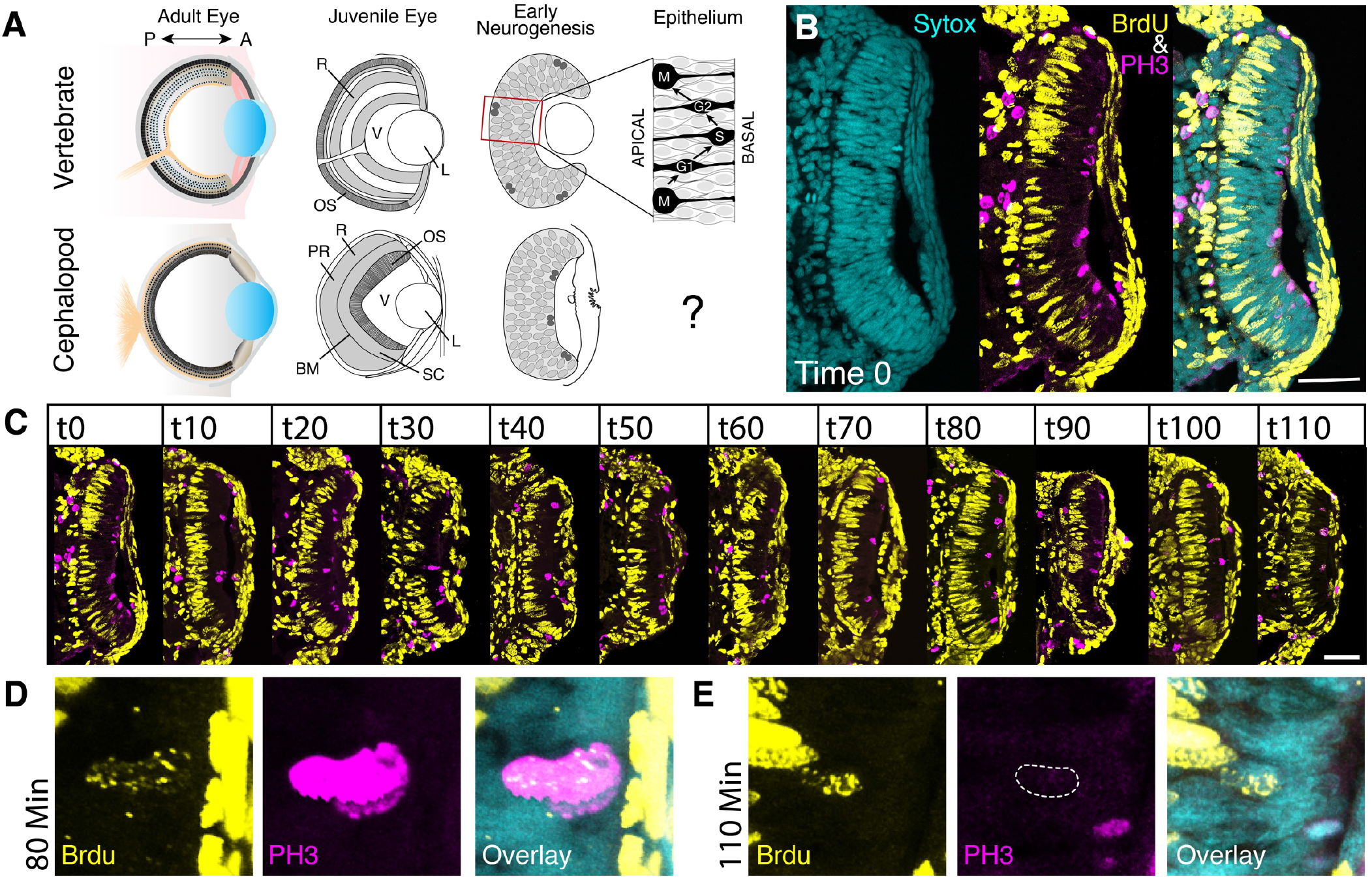
Cell cycle and nuclear migration in the squid retina assessed in fixed tissue. **A.** Schematic of the vertebrate and cephalopod eye at adult, juvenile and early stage development. Box shows enlargement of the vertebrate pseudostratified epithelium, apical to the left. Grey ovals are the nuclei distributed across the tissue early in development, darker grey nuclei are undergoing mitosis. *R:* Retina; *L:* Lens; *OS:* Outer Segment; *V:* Vitreous Space; PR: Photoreceptor Nuclear Layer; *SC:* Support Cell Layer; *BM:* Basal Membrane. **B.** 10 minute BrdU pulse and immediate fix (Time 0) BrdU/PH3 antibody stain shows BrdU incorporation on the posterior side of the epithelium (S-phase) and PH3 stain on the anterior side of the epithelium (M-phase). Scale 50um. **C.** BrdU/PH3 time course. 10 minute BrdU pulse, fixed every 10 minutes followed by BrdU/PH3 antibody stains. Scale 50um. Cell mixing is not observed until 80 minutes after pulse. **D.** 80 minutes after BrdU pulse is the first time point where BrdU+/PH3+ nuclei are found on the apical side of the retina. **E.** 110 minutes after pulse is the first time point where BrdU+/PH3− nuclei are found on the apical side of the retina.

Previous observations from fixed tissue suggested that the cephalopod retinal primordium is a pseudostratified epithelium, which is unusual for an invertebrate neurogenic tissue (Koenig et al, 2016; (Koenig & Gross, 2020) (Figure 1A). This type of neuroepithelium is characteristic of central nervous system development in vertebrates, including the retina (Taverna et al., 2014). A pseudostratified epithelium is a monolayer composed of elongated cells with nuclei distributed along the apicobasal axis. Cells in pseudostratified epithelia undergo nuclear movement correlated with the cell cycle called interkinetic nuclear migration (Norden, 2017). This migration is required for proper differentiation and organization in the developing vertebrate nervous system (Strzyz et al., 2016). During nuclear migration, mitosis occurs on the apical surface and in vertebrates this corresponds to the posterior side of the developing retina (Baye & Link, 2008) (Figure 1A). In the cephalopod, fixed time point data suggests that mitosis occurs on the anterior side of the developing retina, inverted relative to vertebrates (Koenig et al., 2016) (Figure 1A). These observations suggest that neurogenesis may be more similar to vertebrate processes than previously observed in other invertebrates and lead us to ask whether cephalopod retinal progenitor cells undergo interkinetic nuclear migration and how cells differentiate within this epithelium.

## Results

### Cell cycle in the developing squid retina

Previous work in the squid retina showed that at Arnold stage 23, a three hour BrdU pulse resulted in incorporation throughout the retina (Arnold, 1965; Koenig et al., 2016). This long pulse provided time for cells to move after incorporation, making it difficult to determine if only a subset of cells may be proliferative. To more precisely understand cell cycle state and organization in the early retina, we performed a 10 minute BrdU pulse experiment, fixing immediately, and co-labeling with phosphohistone H3 immunofluorescence (PH3) (Figure 1B). We see that BrdU incorporation in S-phase cells is restricted to the posterior of the retina after 10 minutes. We also find PH3 positive labeled nuclei, in late G2 and M phase, on the anterior surface of the retina, supporting previous observations. The lack of BrdU incorporation in the anterior retina suggests that some cells are already post-mitotic.

If cells are undergoing interkinetic nuclear migration, we would expect BrdU positive nuclei to eventually migrate apically to divide. To test this hypothesis, we chased our 10 minute BrdU pulse, sampling embryos every ten minutes for 110 minutes, again co-labeling with PH3 (Figure 1C). We found that at 80 minutes after the pulse we see BrdU positive/PH3 positive cells in the anterior retina, suggesting that nuclei are migrating apically to divide (Figure 1D). If proliferative cells in the posterior retina eventually exit the cell cycle and join the cells in the anterior retina, we would expect to find BrdU positive/PH3 negative cells in the anterior retina. Indeed, at 110 minutes we find BrdU positive, PH3 negative cells in the anterior retina (Figure 1E). Together, these data suggest that cells in the cell cycle have nuclei residing in the posterior retina and these nuclei migrate apically to divide. Eventually these cells exit the cell cycle, and their cell bodies migrate to the anterior of the retina. These data are consistent with cells undergoing interkinetic nuclear migration, with post-mitotic cells segregated apically.

### *In vivo* observations of nuclear migration and proliferation in the squid retinal epithelium

While fixed tissue experiments support our hypothesis, to fully understand the dynamics of these cell behaviors, it is necessary to observe development *in vivo*. This required us to develop a cell-resolution live-imaging protocol in cephalopods (Figure 2A). We injected fluorescent Dextran to broadly label cell membranes and performed >9 hour developmental time course imaging experiments of the squid retina at stage 23 (Figure 2A). Within these time course datasets, we were able to track individual cells migrating from the basal lamina to the apical side of the retina, where they mitose, and migrate basally again (Figure 2B, 2C, Movie S1, S2, S3). These data confirm the process of interkinetic nuclear migration in the squid retina. From these data we are able to calculate nuclear velocity and were able to assess that this migratory behavior is similar in velocity to movement described in the vertebrate retina (Figure 2D) (Norden et al., 2009).

**Figure 2:**
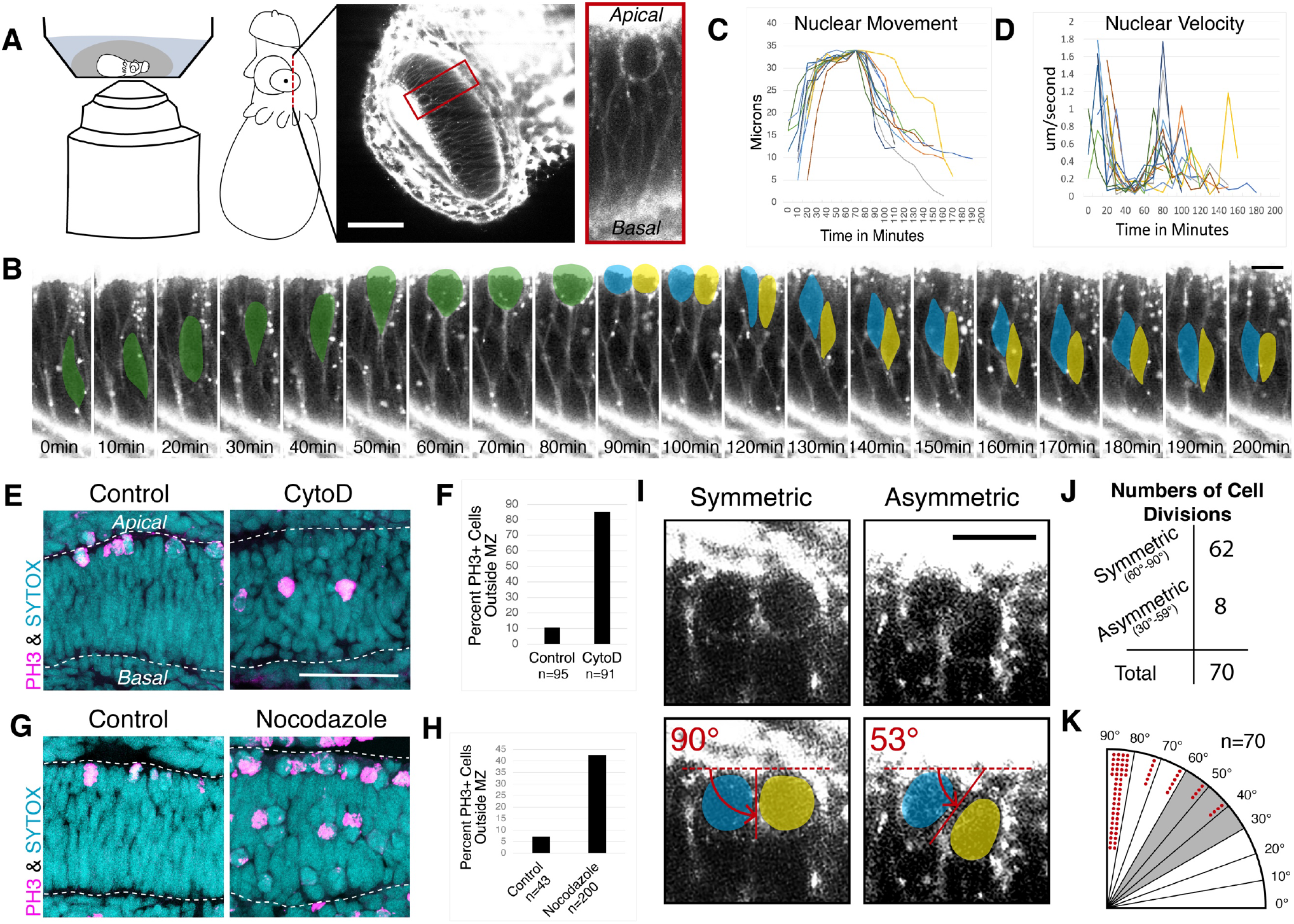
Nuclear migration and mitosis in the retinal epithelium. **A.** Schematic of squid embryo live-imaging. Scale is 50um **B.** False-colored timelapse of a single mitosis in the squid retina at stage 23 showing apical migration in the epithelium, mitosis, and basal migration. Apical is up, basal is down. Scale is 10um. **C.** Graph of nuclear movement as tracked using the Trackmate Fiji plugin (Schindelin et al., 2012). 13 nuclei tracked in 3 embryos. Tracks aligned to the highest point in migration. **D.** Graph of nuclear velocity calculated from the cells tracked in C. **E.** PH3 immunohistochemistry on actin polymerization inhibited squid retina at stage 23. 5uM Cytochalasin D and DMSO control treatment for 7 hours and immediate fix. Scale is 50um. **F.** Quantification of PH3+ nuclei outside the mitotic zone. **G.** PH3 immunohistochemistry on microtubule polymerization inhibited squid retina at stage 23. 5uM Nocodazole and DMSO control treatment for 7 hours and immediate fix. **H.** Quantification of PH3+ nuclei outside of the mitotic zone. **I.** Example of symmetric and asymmetric cell divisions observed in live-imaging experiments. Measurement of angle of division shown. Scale is 10um **J.** Quantification of symmetric (60°-90°) and asymmetric (30°-60°) cell divisions quantified from videos of 5 embryos and a total of 70 mitoses. **K.** Radial histogram quantification of division angles. Each dot represents a single mitosis.

The molecular motors contributing to nuclear migration have been interrogated in a number of developmental contexts and some tissues rely exclusively or primarily on actomyosin, while others require microtubules contribution to promote migration (Kosodo, 2012; E. J. Meyer et al., 2011; Norden, 2017). To understand the contribution of these motors to nuclear migration in the squid retina, we treated stage 23 embryos with either the actin polymerization inhibitor cytochalasin D or the microtubule polymerization inhibitor nocodazole for seven hours and fixed immediately. The impact of these inhibitors on nuclear migration was assessed using immunofluorescence for PH3 (Figure 2E-2H). We find that nuclei positive for PH3 in the cytochalasin D treated retinas are often displaced from the mitotic zone (MZ), defined as the region of the epithelium containing nuclei abutting the most apical membrane. We find a small percentage of PH3 positive nuclei in controls are also outside the mitotic zone, which are likely at the end of G2, migrating toward the apical surface. Quantifying these differences, we see a significant increase in cells outside the mitotic zone in our CytoD treated embryos suggesting actin polymerization is required for nuclear migration. In addition, we see defects in migration in nocodazole treated embryos; a population of PH3 positive nuclei accumulated at the apical surface of the retina. This is what we would expect if microtubules were not required for migration, as cells that have successfully migrated apically would then arrest and accumulate in M phase as microtubules are required for mitosis. However, in addition to this accumulation, we find a subpopulation of PH3 positive cells away from the mitotic zone, suggesting that microtubules play a role in nuclear migration in the cephalopod retina.

In addition to nuclear movement, we were also able to assess the angle of cell division in the epithelium. Symmetrical and asymmetrical cell division is an essential aspect of regulating self-renewal, cell cycle exit and cell fate commitment during neurogenesis in multiple organisms (Yu et al., 2006; Zigman et al., 2005); (Dehay & Kennedy, 2007). Overwhelmingly, we find most mitoses observed to be symmetrical relative to the apical surface of the epithelium (Figure 2I, Movie S2). We hypothesize that these divisions are self-renewing. We also observe approximately 10% of cell divisions to be angled relative to the apical surface (Figure 2I, 2J & 2K, Movie S3). We determined that symmetrical cell divisions had a division plane with an angle relative to the apical surface between 60-90 degrees and asymmetrical divisions between 30-60 degrees (Figure 2K). We did not observe any cell divisions perpendicular to the apical surface, (0-30 degrees) as have been observed in other systems (Knoblich, 2008; Morin & Bellaïche, 2011). We hypothesize that these asymmetric cell divisions may contribute to cells exiting the cell cycle, however we were unable to follow cells long enough during live imaging to confirm this hypothesis.

### Molecular identity during retinal neurogenesis

To better understand the molecular state correlated with the cell cycle organization observed in our BrdU and live imaging experiments, we sought to identify molecular markers that help define differentiation trajectories in the cephalopod retina. We analyzed spatiotemporal expression of thirteen candidate neurogenesis markers in the retina (Figure 3A, Supplemental Figure 1, Supplemental Figure 2). We identified that in early retinal development, stage 21, *DpSoxB1* is homogeneously expressed across the retinal epithelium. At stage 23 the retina is divided across the apical-basal axis with *DpSoxB1* found on the basal side of the retina, correlating with BrdU incorporation, and *DpEphR* is expressed on the apical side. At stage 25 we see the maintenance of segregated *DpSoxB1* and *DpEphR* expression, as well as the first evidence of the terminal differentiation markers *DpRetinochrome* and *DpRhodopsin*. At stage 27, the first evidence of the basal membrane is apparent, which is correlated with a loss of strict segregation of *DpSoxB1* and *DpEphR* expression. We hypothesize this change is a result of cell-body migration or mixing, with apical cells moving to the posterior of the retina and basal cells moving to the anterior of the retina. We hypothesize that this resorting moves the post-mitotic cells to the posterior where they will become the first photoreceptor cell bodies to migrate behind the basal membrane. This is supported by previous observations of nuclei crossing the basal membrane (Koenig et al., 2016). Supporting this hypothesis, we find that *DpSoxB1* expression is never found posterior to the basal membrane. However, *DpEphR* is found on both sides of the membrane until hatching. At hatching *DpEphR* is isolated to a subset of cells posterior to the basal membrane. These cells may be maturing photoreceptor cells that have recently migrated to the photoreceptor cell layer during hatching and post-hatching stage growth. Terminal differentiation markers span both sides of the basal membrane at hatching. We hypothesize that *DpRhodopsin* is expressed exclusively in the photoreceptor cells and is being trafficked to the outer segment and that *DpRetinochrome* is expressed in both the photoreceptor cells and support cells.

**Figure 3:**
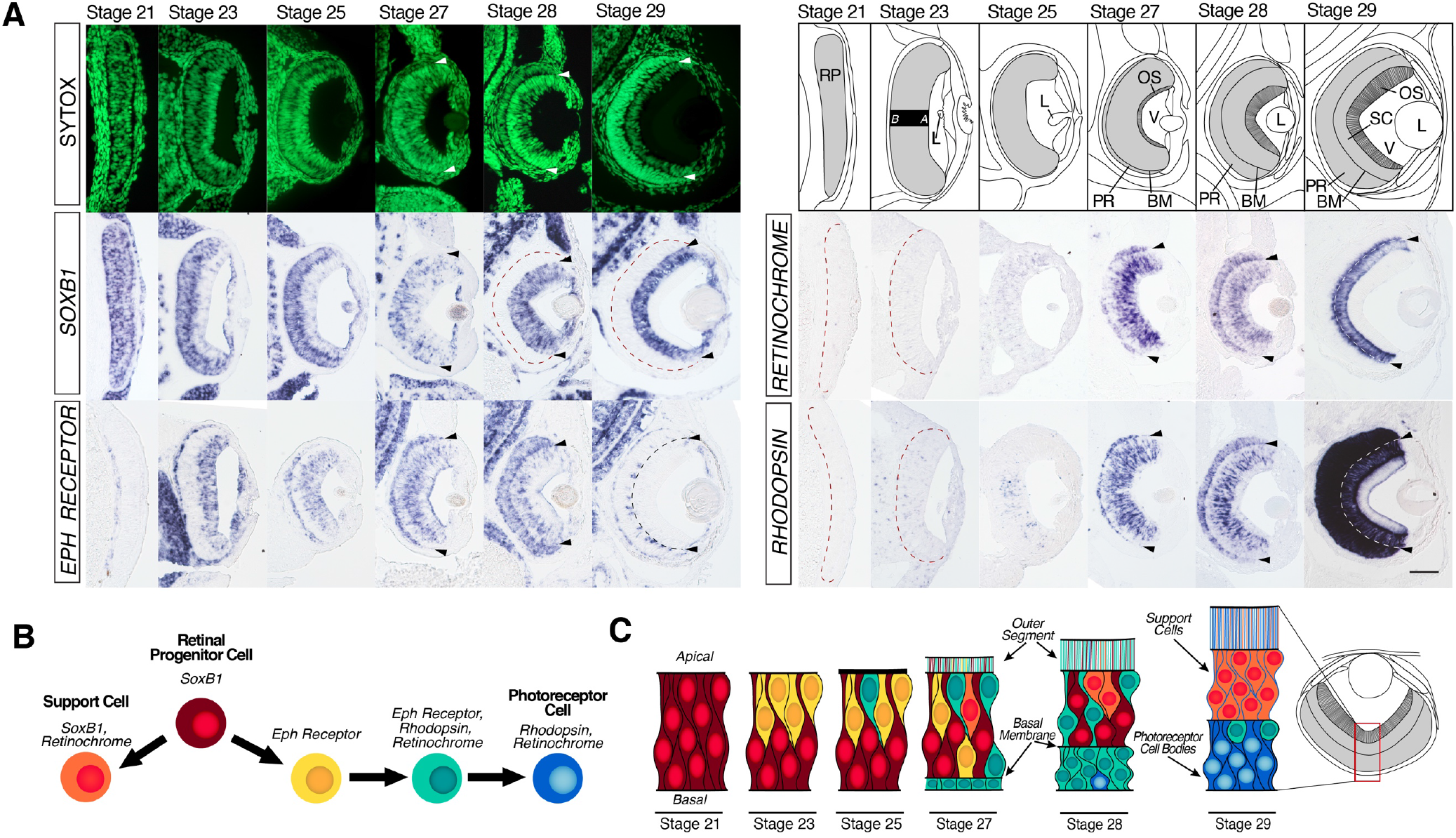
Molecular marker expression suggests a retinal differentiation trajectory. **A.** Staged mRNA expression in the developing cephalopod retina: *DpSoxB1, DpEphR, DpRhodopsin, DpRetinochrome*. Nuclear stain (sytox) shown in green. Schematic of stages labeled and shown. Arrowheads and black and white dotted lines identify the basal membrane. Red dotted lines identify the posterior retinal boundary. Scale is 50um. *RP,* Retinal Placode; *L* Lens; *OS,* Outer Segment; *BM,* Basal Membrane; *PR,* Photoreceptor Cells; *V,* Vitreous Space; *A,* Apical; *B,* Basal. Stage 21, 23, & 25 embryo anterior is down. Stage 27, 28, & 29 embryo dorsal is up. **B.** Hypothesis of cell differentiation trajectories. Color key is used in C. **C.** High magnification summary of time course gene expression data. At stage 21 the retina is homogenous. At stage 23 the retina is divided into apical and basal regions. At stage 25, evidence of terminal differentiation markers, *DpRhodopsin* and *DpRetinochrome*, are evident. At stage 27, the first nuclei are found behind the basal membrane and strict segregation of apical and basal expression is lost. Stage 28, the number of photoreceptor cells behind the basal membrane has increased and gene expression shows increased regional segregation. Stage 29, differentiation markers are robustly expressed.

This candidate gene survey suggests cell differentiation trajectories in the retina (Figure 3B, 3C). Our photoreceptor cell differentiation trajectory starts as *DpSoxB1* positive retinal progenitor cells undergoing interkinetic nuclear migration in the pseudostratified epithelium. These cells transition to *DpEphR* positive cells and become post-mitotic. We hypothesize that this is the result of an asymmetric cell division. Eventually *DpEphR* expressing cells mature and begin to express terminal differentiation markers, ultimately completing differentiation when they exclusively express *DpRhodopsin* and *DpRetinochrome*.

We hypothesize a simple differentiation trajectory for the proliferative support cell population (Figure 3B & 3C). These cells start as *DpSoxB1* positive retinal progenitor cells, undergoing interkinetic nuclear migration. This self-renewing population is maintained throughout development. During cell mixing at stage 27, *DpSoxB1* positive support cells begin to express *DpRetinochrome,* migrate anterior in the retina and remain in the cell cycle. The *DpSoxB1/DpRetinochrome* expressing support cells may be a long term stem cell population in the retina.

### Notch regulates retinal progenitor identity in the cephalopod

Our data show that BrdU incorporation correlates with *DpSoxB1* expression and *DpEphR* expression correlates with post mitotic cells. Our BrdU chase experiments suggest that *DpSoxB1* cells likely become *DpEphR* cells over time (Figure 4A, 4B & 4C). Notch signaling is a well-known regulator of cell cycle exit and differentiation. Previous work showed that inhibiting Notch signaling in the cephalopod using the gamma-secretase inhibitor DAPT led to a loss of BrdU incorporation in the retina, showing that Notch was inhibiting cell cycle exit. We were interested in understanding the role of Notch signaling in regulating cell fate trajectories in conjunction with proliferation. We first performed *in situ* hybridization for Notch signaling pathway members in the retina (Figure 4D and Supplemental Figure 3). If Notch signaling is important for maintaining progenitor fate, we would expect Notch and Hes to be enriched in retinal progenitor cells. We find that *DpNotch* and *DpHes-1* expression is correlated with *DpSoxB1* expression and retinal progenitor identity (Figure 4D and Supplemental Figure 3). Interestingly, we find that Delta is ubiquitously expressed during these stages (Supplemental Figure 3). We also find *DpNotch* expressed in a limited number of nuclei on the apical side of the retina. We wanted to know if this expression correlated with nuclei that had migrated apically to divide. We performed *in situ* hybridization for *DpNotch* in conjunction with PH3 immunofluorescence and found that apical *DpNotch* expression is correlated with PH3 positive nuclei (Figure 4E). Inheritance of Notch mRNA may be an important part of maintaining progenitor cell identity.

**Figure 4:**
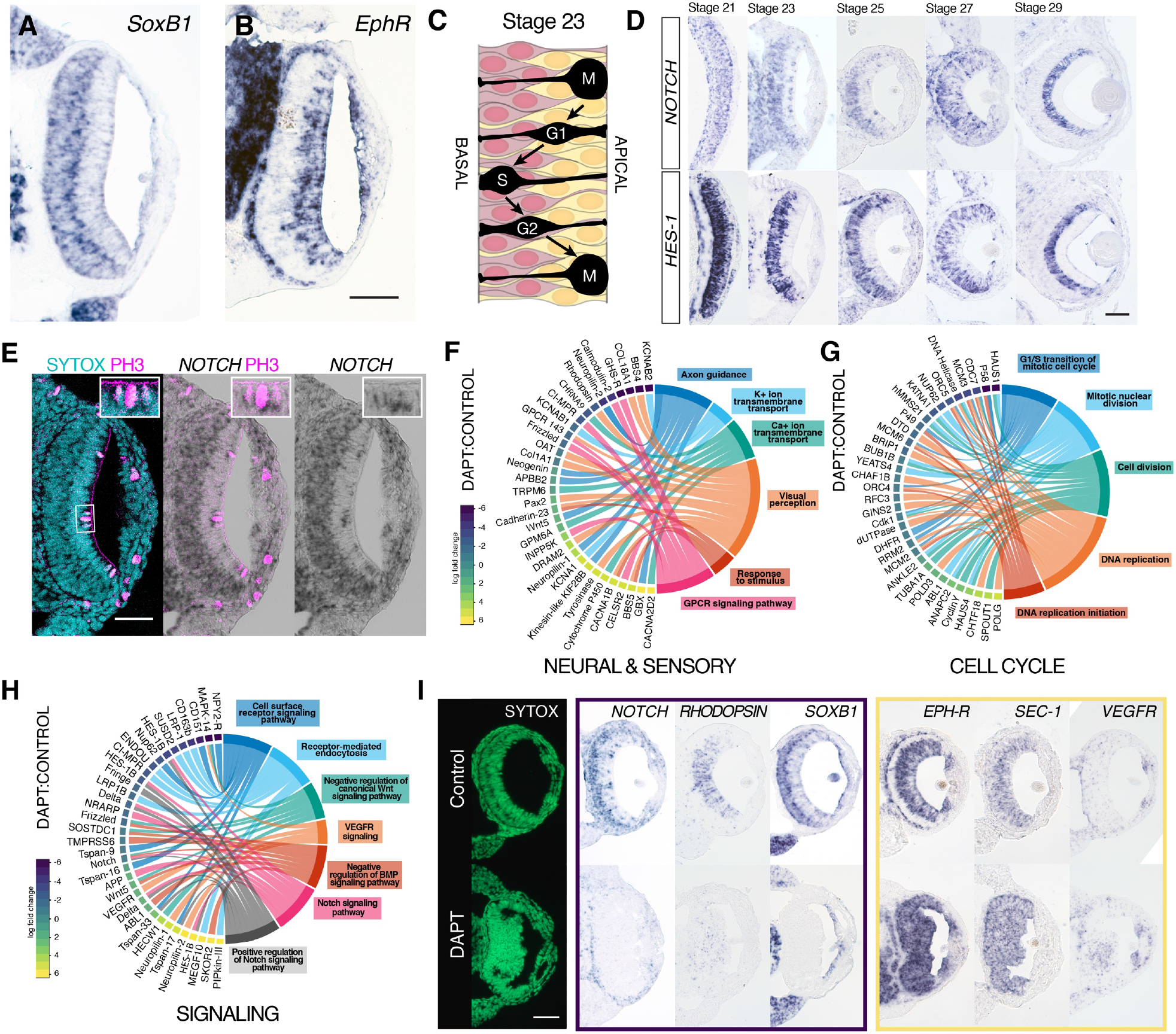
Notch signaling is required to maintain retinal pr-ogenitor cell identity. **A.** *DpSoxB1* expression on the basal side of the retina at stage 23 in retinal progenitor cells. **B.** *DpEphR* expression on the apical side of the retina at stage 23 in post-mitotic cells. Scale 50um. **C.** Schematic of nuclear migration within the pseudostratified retinal epithelium. Red (*DpSoxB1*) and yellow (*DpEphR*) color corresponds to gene expression and summary Figure 3B and 3C. **D.** *DpNotch* and *DpHes-1* expression correlates with *DpSoxB1* expression in the retinal progenitor cell population. Scale 50um **E.** *DpNotch* expression on the apical side of the retinal epithelium correlates with PH3 expression. Scale 50um. **F.** DAVID GO-annotations of RNA-seq results of Notch inhibited eyes. Embryos treated with 20uM DAPT and DMSO control at stage 23 for 24 hours. Eyes dissected for sequencing. All results shown have a p-value < 0.05. Neural sensory genes **G.** Cell-cycle genes **H.** Signaling pathway genes. Color Key is shared for all RNA-seq results. **I.** *In situ* hybridization on Control and Notch inhibited retina. Embryos treated with 20uM DAPT and DMSO control at stage 23 for 24 hours and fixed immediately. Nuclear stain (SYTOX-green) shown in green in the first column. Selected down-regulated genes in DAPT embryos in the purple box on the left. Selected up-regulated genes in DAPT embryos in the yellow box on the right. *DpSoxB1* expression, the retinal progenitor marker, is lost from the retina in Notch inhibited embryos. *DpEphR* expression, the neuroblast marker, has expanded expression throughout the retina in Notch inhibited embryos.

To assess the role of Notch in cell fate we used the gamma secretase inhibitor DAPT to inhibit active signaling. Embryos were treated with DAPT or DMSO starting at stage 23 for 24 hours. Eyes were dissected and pooled. RNA was extracted from three experimental and control samples and sequenced. 2242 genes were differentially expressed between DAPT and control eyes with a p-value of less than 0.05 (Supplemental Table 1). *DpSoxB1* was found downregulated and *DpEphR* was found upregulated in the dataset, but these changes were not deemed statistically significant. DAPT compared to control eye samples showed both up and down regulation in neural and sensory markers. Of note is the down regulation of Rhodopsin and other G-protein coupled receptor related genes (Figure 4F). In addition, cell cycle related genes were downregulated, which is expected as DAPT treatment leads to cell cycle exit in the squid retina (Koenig et al., 2016) (Figure 4G). Finally, DAPT versus control eyes showed significant changes in cell signaling genes, suggesting that Notch may self-regulate as well as interact with other signaling pathways in the retina, including VEGFR, BMP and Wnt (Figure 4H).

To confirm changes in gene expression identified in the RNA-seq analysis, we performed *in situ* hybridization studies on control and DAPT embryos (Figure 4I and Supplemental Figure 4 and Supplemental Figure 5). We confirm trends shown in the RNA-seq results. Specifically, we find a loss of *DpNotch* expression, a downregulation of *DpRhodopsin,* and a complete loss of *DpSoxB1* in DAPT treated retinas (Figure 4I). We also observe a gain of expression of *DpEphR* and synaptic transmission related gene, *DpSec-1,* in the posterior retina (Figure 4I). Interestingly, we see ectopic expression of *DpVEGFR* in a subpopulation of cells in the DAPT treated retinas relative to control. The unusual placement of ectopic expression of *DpVEGFR* may be compounded by the disorganization of the DAPT treated retinas. The complete loss of *DpSoxB1* and the ectopic expression of *DpEphR* shows that DAPT inhibition not only results in changes in cell cycle state, but also changes molecular fate in the squid retina. These data suggest that Notch signaling is required to maintain retinal progenitor identity in the squid.

## Discussion

In the current study we have characterized the nuclear oscillatory behaviors of retinal progenitor cells in squid. We have defined the molecular markers for progenitor, post-mitotic, and fully differentiated cells in the retina and we have shown that Notch signaling is regulating both cell cycle and molecular identity during retinal development. The importance of these findings is in comparison to what is known about neurogenesis in other systems.

Pseudostratified epithelia and interkinetic nuclear migration have been observed in multiple tissue types and across multiple species (E. J. Meyer et al., 2011). In the nervous system, pseudostratification was historically considered a vertebrate specific developmental trait, which was responsible for the large size of the central nervous system. Elaborate tissue-level morphogenesis is has not been observed in invertebrate neural development, where individual cells delaminate or ingress from a simple or stratified neuroepithelium, sometimes divide, and then migrate to their final destination to differentiate (N. P. Meyer & Seaver, 2009; Sur et al., 2020); (Byrne, 2019; Hartenstein & Stollewerk, 2015). However, interkinetic nuclear migration has been described in the *Drosophila* optic lobe (Egger et al., 2007). In this case, the apicobasal distance of migration is significantly smaller than vertebrate neurogenic tissues and cells delaminate from the edge of the proliferative epithelium and migrate to their final destination to differentiate (Egger et al., 2007; Holguera & Desplan, 2018; Strzyz et al., 2016). This process has been compared to “conveyor belt” neurogenesis found in the zebrafish ciliary and tectal marginal zones, where, during long term growth, cells are progressively added from the edge of a pseudostratified stem cell population (Joly et al., 2016). The behavior described in the squid retina is unusual because of its unique similarity to early neurogenesis in vertebrate species, which is characterized by waves of differentiation within the pseudostratified epithelium (Agathocleous & Harris, 2009; Livesey & Cepko, 2001). In the squid, we show that apicobasal height of the epithelium is similar to vertebrate tissues, cells undergo interkinetic nuclear migration while proliferative, and cells remain within the bounds of the epithelium after delaminating and differentiating. The functional difference between conveyor belt neurogenesis versus within-epithelium differentiation is unclear, but the architecture of the epithelium may provide physical landmarks or signaling information important for proper organization.

Although pseudostratified neurogenesis is found in the cephalopod retina, the cell biology of chordal or central nervous system neurogenesis is still not well described. It has been reported that cells migrate from the lateral lip tissue, an embryonic neurogenic tissue found in the cephalopod embryo to regions within the brain but pseudostratification of the lateral lips is not apparent (Deryckere et al., 2021). Cephalopod central nervous system development may more closely resemble neurogenesis found in other invertebrates or molluscs (Deryckere et al., 2021; Marthy, 1987). With a central nervous system composed of significantly more cells than other Spiralians and requiring long distance cell migration, it is likely that cephalopods have evolved lineage specific mechanisms to manage this process.

Interestingly, in the squid, candidate gene expression of neurogenic factors also differs significantly between the developing retina and other brain regions. This includes a lack of expression of the canonical neurogenic ELAV, *DpELAV*, in the retina, an RNA binding protein commonly found in differentiating neurons across species (Akamatsu et al., 2005; Denes et al., 2007; N. P. Meyer & Seaver, 2009; Nakanishi et al., 2012; Nomaksteinsky et al., 2009). bHLH factors *DpNeuroD* and *DpNeuroG*, both with orthologs that have roles in neural specification and differentiation in many animals, are expressed in the lateral lips and brain in *D. pealeii* and other cephalopods but are completely absent for the retina (Supplemental Figure 2) (Buresi et al., 2013; Deryckere et al., 2021; Shigeno et al., 2015). However, it has been reported in *Octopus vulgaris* that achaete-scute homolog *Ov-ascl1*, a third proneural bHLH transcription factor, is expressed in the retina (Deryckere et al., 2021). Furthermore, SoxB1, required to maintain neural precursors cells in vertebrates (Sox1, 2 and 3) and neuroblasts in *Drosophila* (SoxNeuro and Dichaete), is also a retinal progenitor marker in cephalopods (Buescher et al., 2002; Bylund et al., 2003; Ferrero et al., 2014; Holmberg et al., 2008; Overton et al., 2002; Taranova et al., 2006). We also find that both cephalopod-specific *DpELAVL* paralogs show expression that suggests a role in neurogenesis in both the retina and brain (Supplemental Figure 2).

Finally, we show that Notch signaling is maintaining molecular identity of the progenitor population in the squid retina, and that the loss of Notch signaling leads to expression of the post-mitotic marker *DpEphR*. In the vertebrate retina, the premature loss of Notch signaling leads to the differentiation of early born cell types (Perron & Harris, 2000). In *D. pealeii*, we see a decrease in *DpRhodopsin* expression, suggesting cells may have arrested in this immature post-mitotic state. We also see the maintenance in *DpRetinochrome* expression, which may be evidence of differing trajectories of these cell types during neurogenesis.

The function of pseudostratification and interkinetic nuclear migration during development remains unclear, however our data sheds new light on old hypotheses. In particular, pseudostratification may aid in intracellular organization, as we see significant mRNA localization in our data, including mRNA transport correlated with nuclear migration. It may also be that the physics and dynamics of pseudostratified epithelia are better at accommodating increased proliferation or provides an organizational scaffold for the differentiating network of neurons. It will be interesting to apply new fluid dynamics approaches used to study vertebrate neurogenesis to the cell biology of this similar tissue (Azizi et al., 2020). Ultimately, the evidence we have generated in cephalopods suggests that vertebrate-like cell behaviors during neurogenesis are not exclusive to the chordate lineage. With greater sampling we may better understand the evolutionary changes that contribute to the diversity and complexity we see.

## Methods

### Animal Husbandry

*Doryteuthis pealeii* egg sacks were obtained from the Marine Biological Labs. Egg sacks were kept at 20 degrees Celsius in 20 gallon aquaria in artificial seawater under a day/night cycle. Although not required, European guidelines for cephalopod research were followed.

### Cloning and *in situ* Hybridization

Primers were designed using Primer3 in the Geneious software package version 2020.04 (https://www.geneious.com) and primer sequences are reported in Supplemental Table 2. Genes were cloned into pGEM-T Easy vector and confirmed with Sanger sequencing and DIG-labeled RNA probes were synthesized as previously reported (Koenig et al., 2016). Embryos were fixed overnight at 4 degrees Celsius, washed and dehydrated stepwise into 100% ethanol. Embryos were embedded, paraffin sectioned and *in situ* hybridization was performed as previously reported (Neal et al., n.d.). All *in situs* were replicated in at least three embryos, across multiple separate *in situ* experiments. Slides were stained overnight with Sytox-Green 1:1000 overnight, mounted using ImmunoHistoMount (Abcam) and imaged on a Zeiss Axioskop 2.

### BrdU Experiments

Embryos were bathed in BrdU (10 mM) in pen-strep seawater for 10 minutes as previously described (Koenig et al., 2016). Embryos were fixed immediately, and the remaining embryos were moved into pen-strep seawater. Groups of 10-15 embryos were fixed every ten minutes for two hours. Embryos were fixed overnight at 4 degrees and washed out into PBS-Tween. Embryos were stepped into 30% sucrose and embedded in tissue freezing medium and sectioned as previously described (Koenig et al., 2016). Antigen-retrieval and immunofluorescence was performed as previously described. BrdU antibody (Abcam ab6326) at concentration 1:250. Phosphohistone H3 antibody (Sigma-Aldrich 06-570) was used at concentration 1:300. Secondary antibodies goat anti-rat Alexa Fluor 488 and goat anti rabbit Alexa Fluor 647 (Invitrogen). Sections were counterstained with SYTOX-Green at 1:1000. Sections were imaged on a Zeiss LSM 880 or Zeiss LSM 980.

### Live-imaging experiments

Embryos were dissected from egg cases. The vitreous space was injected using a pico-liter microinjector with Dextran Alexa Fluor 488 10,000 MW (D22910) at stage 23. Embryos were embedded in 1% low melt agarose in seawater and mounted in cover glass bottom dishes (100503-366). Embryos were immersed in pen strep seawater. Embryos were imaged on a Zeiss 880. Embryos were imaged every ten minutes for at least nine hours.

### Imaging analysis

Image analysis was performed in Fiji (Schindelin et al., 2012). Intensity range was adjusted in Fiji to better identify cell membranes. Drift correction was performed using Fiji plugin CoordinateShift (written by Housei Wada, https://signaling.riken.jp/en/en-tools/imagej/). Nuclear tracking was performed both manually and using Fiji plugin Trackmate (Robertson, 2018; Schindelin et al., 2012). Tracks were visualized and distance and velocity measurements were obtained from Trackmate and plotted graphically, normalizing to the highest point in migration.

### *Ex ovo* and Drug Treatments

Embryos were treated in 20uM DAPT solution or DMSO control in filter-sterilized, Pen-Strep seawater starting at stage 23 for 24 hours and fixed immediately as previously described (Koenig et. al, 2016). Embryos were bathed in 5uM Nocodazole (M1404-2MG, Sigma-Aldrich) and 5uM Cytochalasin D (C8273-1MG, Sigma-Aldrich) for 7 hours and fixed immediately.

### RNA-seq and Bioinformatics

Stage 23 embryos were treated with the gamma-secretase inhibitor DAPT at 20um in filter sterilized sea water for 24 hours and Control embryos were treated with the equivalent amount of DMSO, as previously described (Koenig et al., 2016). DAPT and Control eyes were dissected, pooled and macerated in TRIzol and stored at −80 degrees Celsius. RNA was extracted using a standard TRIzol (Invitrogen #15596026) chloroform extraction and passed through a gDNA eliminator mini-spin column (Qiagen #1030958). RNA was precipitated with isopropanol and then again precipitated with ethanol and checked for quality. Library prep and sequencing was performed at the Bauer Core at Harvard University. RNA-seq libraries were generated using the Kapa mRNA-Hyper Prep kit with Poly-A Selection (Roche, Basal) and were sequenced on the Illumina NovaSeq (>70 million 2×150 bp sequences) (Illumina, San Diego, CA).

Sequence quality control was performed according to the best practice recommendation on the Harvard FAS Informatics pipeline (https://informatics.fas.harvard.edu/best-practices-for-de-novo-transcriptome-assembly-with-trinity.html). Erroneous kmers were removed from the paired end Illumina dataset using rCorrector. Reads with Ns or other low complexity pairs were removed using a custom python script provided by the Harvard Informatics GitHub (FilterUncorrectabledPEfastq.py). Adapters and low quality bases were removed using TrimGalore! Reads that mapped using Bowtie2 to the rRNA databases SILVA 132 SSURef Nr99 tax and SILVA 132 LSUParc tax were removed. Pseudomapping was performed by Kallisto with 100 bootstraps to a previously published whole embryo transcriptome (Koenig et al., 2016). Transcript abundances were imported using tximport (https://bioconductor.org/packages/3.7/bioc/vignettes/tximport/inst/doc/tximport.html#use-with-downstream-bioconductor-dge-packages) into DESeq2 using the Kallisto abundance.h5 files. Differential gene expression was determined by importing transcript level abundances and gene level offset using *DESeqDataSetFromTximport* (Soneson et al., 2015). The DESeq2 pipeline was run and differentially expressed genes were considered with a p value of .05 and log2 fold change of above 1 and below −1. All genes shown in the chord plots in figure 4 meet these criteria with the exception of Notch which was included with a log2 fold change of −0.83. GO annotations were identified using the DAVID functional annotation tool (Huang et al., 2007). Chord plots were generated using the R package GOPlot (Walter et al., 2015) and gplots (*Website*, n.d.)Warnes(*Website*, n.d.).

## Supporting information

Supplemental Movie 2 Symmetric

Supplemental Movie 3 Asymmetric

## Acknowledgements

We would like to thank the Srivastava lab for helpful discussions and Mansi Srivastava and Jeffrey Gross for comments on the manuscript. We thank Kevin Woods and the John Harvard Distinguished Science Fellows community for support and the Marine Biological Labs and the Marine Resources Center for assistance and access to embryos.

## Author Contributions

K.M.K conceived of the study with input from F.N., C.M.D. and S.N.; K.M.K, F.N., C.M.D., S.N., A.Z., and A.L. performed experiments. K.J.M. performed phylogenetic analyses. K.M.K., C.M.D and K.J.M. wrote the manuscript in consultation with all authors.

