## Supplemental Figures 1-5 and Supplemental Table 2 for "Cephalopod Retinal Development Shows Vertebrate-like Mechanisms of Neurogenesis"

**Supplemental Figure 1: Phylogenetic analyses for genes identified in this study**

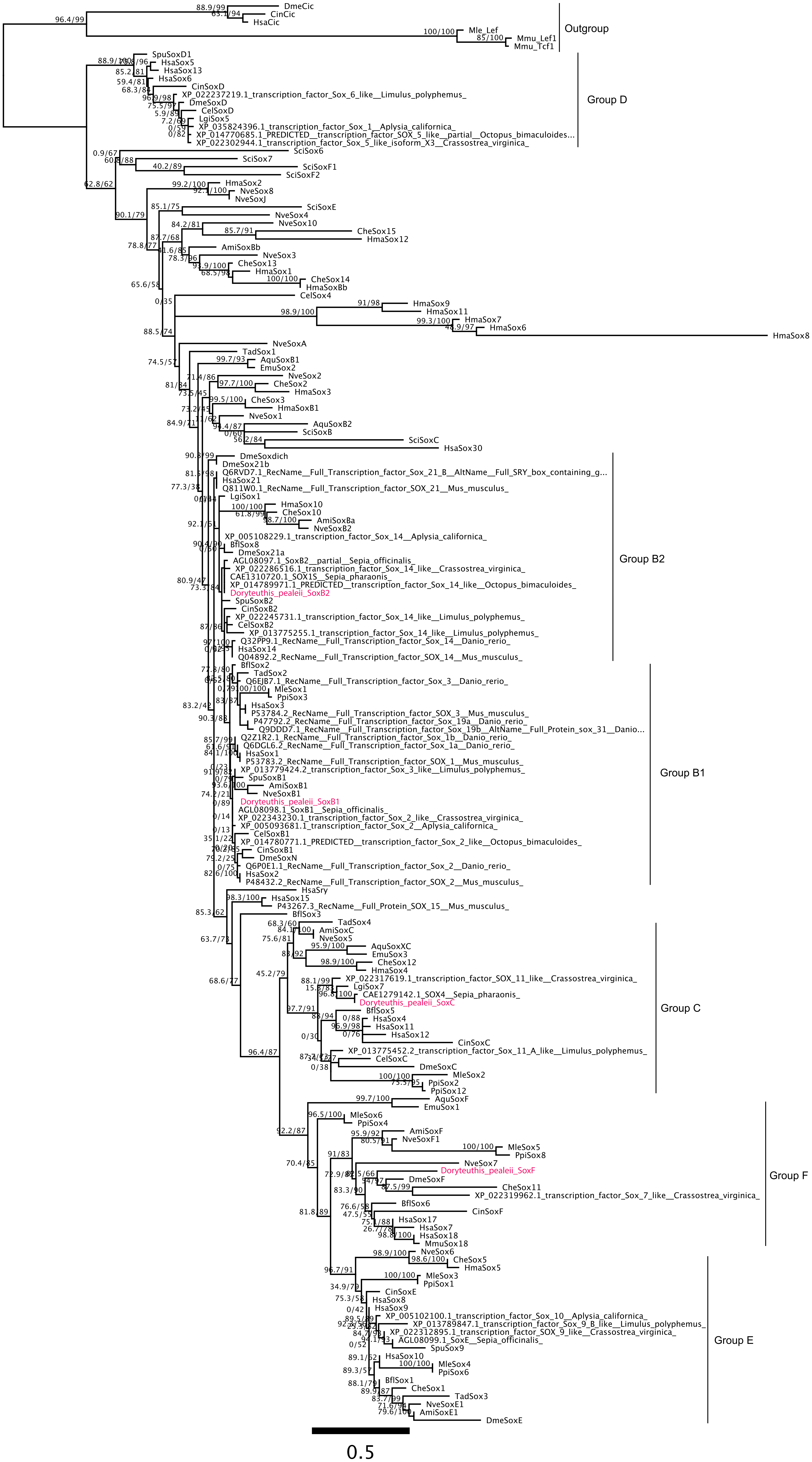

DmeCic
CinCic
HsaCic
Mle_Lef
Mmu_Lef1
Mmu_Tcf1
Outgroup
SpuSoxD1
HsaSox5
HsaSox13
HsaSox6
CinSoxD
XP_022237219.1_transcription_factor_Sox_6_like__Limulus_polyphemus_
DmeSoxD
CelSoxD
LgiSox5
XP_035824396.1_transcription_factor_Sox_1__Aplysia_californica_
XP_014770685.1_PREDICTED__transcription_factor_SOX_5_like__partial__Octopus_bimaculoides...
XP_022302944.1_transcription_factor_Sox_5_like_isoform_X3__Crassostrea_virginica_
Group D
SciSox6
SciSox7
SciSoxF1
SciSoxF2
HmaSox2
NveSox8
NveSoxJ
SciSoxE
NveSox4
NveSox10
CheSox15
HmaSox12
AmiSoxBb
NveSox3
CheSox13
HmaSox1
CheSox14
HmaSoxBb
CelSox4
HmaSox9
HmaSox11
HmaSox7
HmaSox6
HmaSox8
NveSoxA
TadSox1
AquSoxB1
EmuSox2
NveSox2
CheSox2
HmaSox3
CheSox3
HmaSoxB1
NveSox1
AquSoxB2
SciSoxB
SciSoxC
HsaSox30
DmeSoxdich
DmeSox21b
Q6RVD7.1_RecName__Full_Transcription_factor_Sox_21_B__AltName__Full_SRY_box_containing_g...
HsaSox21
Q811W0.1_RecName__Full_Transcription_factor_SOX_21__Mus_musculus_
LgiSox1
HmaSox10
CheSox10
AmiSoxBa
NveSoxB2
XP_005108229.1_transcription_factor_Sox_14__Aplysia_californica_
BflSox8
DmeSox21a
AGL08097.1_SoxB2__partial__Sepia_officinalis_
XP_022286516.1_transcription_factor_Sox_14_like__Crassostrea_virginica_
CAE1310720.1_SOX1S__Sepia_pharaonis_
XP_014789971.1_PREDICTED__transcription_factor_Sox_14_like__Octopus_bimaculoides_
Doryteuthis_pealeii_SoxB2
SpuSoxB2
CinSoxB2
XP_022245731.1_transcription_factor_Sox_14_like__Limulus_polyphemus_
CelSoxB2
XP_013775255.1_transcription_factor_Sox_14_like__Limulus_polyphemus_
Q32PP9.1_RecName__Full_Transcription_factor_Sox_14__Danio_rerio_
HsaSox14
Q04892.2_RecName__Full_Transcription_factor_SOX_14__Mus_musculus_
Group B2
BflSox2
TadSox2
Q6EJB7.1_RecName__Full_Transcription_factor_Sox_3__Danio_rerio_
MleSox1
PpiSox3
HsaSox3
P53784.2_RecName__Full_Transcription_factor_SOX_3__Mus_musculus_
P47792.2_RecName__Full_Transcription_factor_Sox_19a__Danio_rerio_
Q9DDD7.1_RecName__Full_Transcription_factor_Sox_19b__AltName__Full_Protein_sox_31__Danio...
Q2Z1R2.1_RecName__Full_Transcription_factor_Sox_1b__Danio_rerio_
Q6DGL6.2_RecName__Full_Transcription_factor_Sox_1a__Danio_rerio_
HsaSox1
P53783.2_RecName__Full_Transcription_factor_SOX_1__Mus_musculus_
XP_013779424.2_transcription_factor_Sox_3_like__Limulus_polyphemus_
SpuSoxB1
AmiSoxB1
NveSoxB1
Doryteuthis_pealeii_SoxB1
AGL08098.1_SoxB1__Sepia_officinalis_
XP_022343230.1_transcription_factor_Sox_2_like__Crassostrea_virginica_
XP_005093681.1_transcription_factor_Sox_2__Aplysia_californica_
CelSoxB1
XP_014780771.1_PREDICTED__transcription_factor_Sox_2_like__Octopus_bimaculoides_
CinSoxB1
DmeSoxN
Q6P0E1.1_RecName__Full_Transcription_factor_Sox_2__Danio_rerio_
HsaSox2
P48432.2_RecName__Full_Transcription_factor_SOX_2__Mus_musculus_
Group B1
HsaSry
HsaSox15
P43267.3_RecName__Full_Protein_SOX_15__Mus_musculus_
BflSox3
TadSox4
AmiSoxC
NveSox5
AquSoxXC
EmuSox3
CheSox12
HmaSox4
XP_022317619.1_transcription_factor_SOX_11_like__Crassostrea_virginica_
LgiSox7
CAE1279142.1_SOX4__Sepia_pharaonis_
Doryteuthis_pealeii_SoxC
BflSox5
HsaSox4
HsaSox11
HsaSox12
CinSoxC
XP_013775452.2_transcription_factor_Sox_11_A_like__Limulus_polyphemus_
CelSoxC
DmeSoxC
MleSox2
PpiSox2
PpiSox12
Group C
AquSoxF
EmuSox1
MleSox6
PpiSox4
AmiSoxF
NveSoxF1
MleSox5
PpiSox8
NveSox7
Doryteuthis_pealeii_SoxF
DmeSoxF
CheSox11
XP_022319962.1_transcription_factor_Sox_7_like__Crassostrea_virginica_
BflSox6
CinSoxF
HsaSox17
HsaSox7
HsaSox18
MmuSox18
Group F
NveSox6
CheSox5
HmaSox5
MleSox3
PpiSox1
CinSoxE
HsaSox8
HsaSox9
XP_005102100.1_transcription_factor_Sox_10__Aplysia_californica_
XP_013789847.1_transcription_factor_Sox_9_B_like__Limulus_polyphemus_
XP_022312895.1_transcription_factor_SOX_9_like__Crassostrea_virginica_
AGL08099.1_SoxE__Sepia_officinalis_
SpuSox9
HsaSox10
MleSox4
PpiSox6
BflSox1
CheSox1
TadSox3
NveSoxE1
AmiSoxE1
DmeSoxE
Group E
0.5

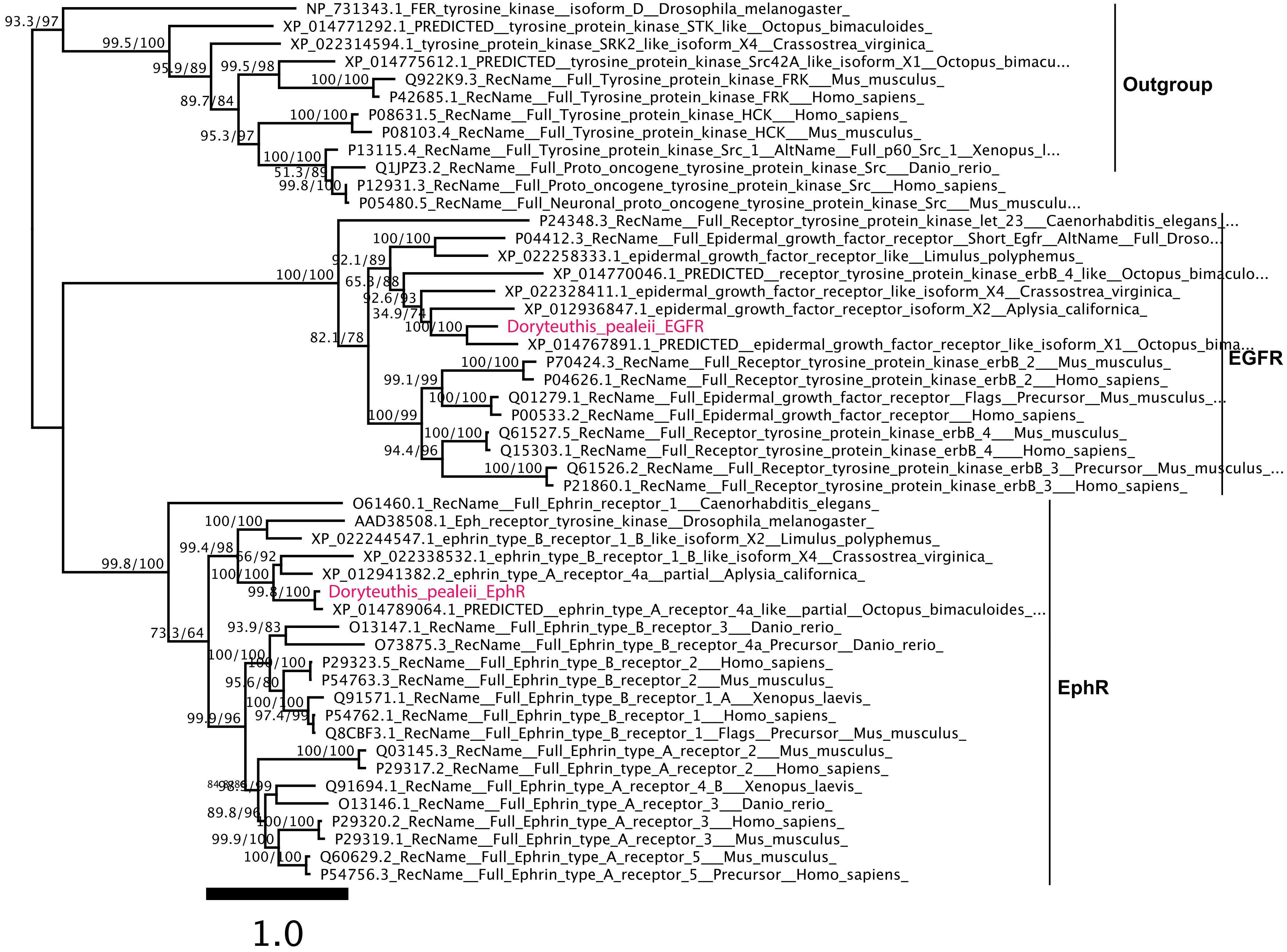
NP_731343.1_FER_tyrosine_kinase__isoform_D__Drosophila_melanogaster_
XP_014771292.1_PREDICTED__tyrosine_protein_kinase_STK_like__Octopus_bimaculoides_
XP_022314594.1_tyrosine_protein_kinase_SRK2_like_isoform_X4__Crassostrea_virginica_
XP_014775612.1_PREDICTED__tyrosine_protein_kinase_Src42A_like_isoform_X1__Octopus_bimacu...
Q922K9.3_RecName__Full_Tyrosine_protein_kinase_FRK___Mus_musculus_
P42685.1_RecName__Full_Tyrosine_protein_kinase_FRK___Homo_sapiens_
P08631.5_RecName__Full_Tyrosine_protein_kinase_HCK___Homo_sapiens_
P08103.4_RecName__Full_Tyrosine_protein_kinase_HCK___Mus_musculus_
P13115.4_RecName__Full_Tyrosine_protein_kinase_Src_1__AltName__Full_p60_Src_1__Xenopus_l...
Q1JPZ3.2_RecName__Full_Proto_oncogene_tyrosine_protein_kinase_Src___Danio_rerio_
P12931.3_RecName__Full_Proto_oncogene_tyrosine_protein_kinase_Src___Homo_sapiens_
P05480.5_RecName__Full_Neuronal_proto_oncogene_tyrosine_protein_kinase_Src___Mus_musculu...
Outgroup
P24348.3_RecName__Full_Receptor_tyrosine_protein_kinase_let_23___Caenorhabditis_elegans_...
P04412.3_RecName__Full_Epidermal_growth_factor_receptor__Short_Egfr__AltName__Full_Droso...
XP_022258333.1_epidermal_growth_factor_receptor_like__Limulus_polyphemus_
XP_014770046.1_PREDICTED__receptor_tyrosine_protein_kinase_erbB_4_like__Octopus_bimaculo...
XP_022328411.1_epidermal_growth_factor_receptor_like_isoform_X4__Crassostrea_virginica_
XP_012936847.1_epidermal_growth_factor_receptor_isoform_X2__Aplysia_californica_
Doryteuthis_pealeii_EGFR
XP_014767891.1_PREDICTED__epidermal_growth_factor_receptor_like_isoform_X1__Octopus_bima...
P70424.3_RecName__Full_Receptor_tyrosine_protein_kinase_erbB_2___Mus_musculus_
P04626.1_RecName__Full_Receptor_tyrosine_protein_kinase_erbB_2___Homo_sapiens_
Q01279.1_RecName__Full_Epidermal_growth_factor_receptor__Flags__Precursor__Mus_musculus_...
P00533.2_RecName__Full_Epidermal_growth_factor_receptor___Homo_sapiens_
Q61527.5_RecName__Full_Receptor_tyrosine_protein_kinase_erbB_4___Mus_musculus_
Q15303.1_RecName__Full_Receptor_tyrosine_protein_kinase_erbB_4___Homo_sapiens_
Q61526.2_RecName__Full_Receptor_tyrosine_protein_kinase_erbB_3__Precursor__Mus_musculus_...
P21860.1_RecName__Full_Receptor_tyrosine_protein_kinase_erbB_3___Homo_sapiens_
EGFR
O61460.1_RecName__Full_Ephrin_receptor_1___Caenorhabditis_elegans_
AAD38508.1_Eph_receptor_tyrosine_kinase__Drosophila_melanogaster_
XP_022244547.1_ephrin_type_B_receptor_1_B_like_isoform_X2__Limulus_polyphemus_
XP_022338532.1_ephrin_type_B_receptor_1_B_like_isoform_X4__Crassostrea_virginica_
XP_012941382.2_ephrin_type_A_receptor_4a__partial__Aplysia_californica_
Doryteuthis_pealeii_EphR
XP_014789064.1_PREDICTED__ephrin_type_A_receptor_4a_like__partial__Octopus_bimaculoides_...
O13147.1_RecName__Full_Ephrin_type_B_receptor_3___Danio_rerio_
O73875.3_RecName__Full_Ephrin_type_B_receptor_4a_Precursor__Danio_rerio_
P29323.5_RecName__Full_Ephrin_type_B_receptor_2___Homo_sapiens_
P54763.3_RecName__Full_Ephrin_type_B_receptor_2___Mus_musculus_
Q91571.1_RecName__Full_Ephrin_type_B_receptor_1_A___Xenopus_laevis_
P54762.1_RecName__Full_Ephrin_type_B_receptor_1___Homo_sapiens_
Q8CBF3.1_RecName__Full_Ephrin_type_B_receptor_1__Flags__Precursor__Mus_musculus_
Q03145.3_RecName__Full_Ephrin_type_A_receptor_2___Mus_musculus_
P29317.2_RecName__Full_Ephrin_type_A_receptor_2___Homo_sapiens_
Q91694.1_RecName__Full_Ephrin_type_A_receptor_4_B___Xenopus_laevis_
O13146.1_RecName__Full_Ephrin_type_A_receptor_3___Danio_rerio_
P29320.2_RecName__Full_Ephrin_type_A_receptor_3___Homo_sapiens_
P29319.1_RecName__Full_Ephrin_type_A_receptor_3___Mus_musculus_
Q60629.2_RecName__Full_Ephrin_type_A_receptor_5___Mus_musculus_
P54756.3_RecName__Full_Ephrin_type_A_receptor_5__Precursor__Homo_sapiens_
EphR
1.0

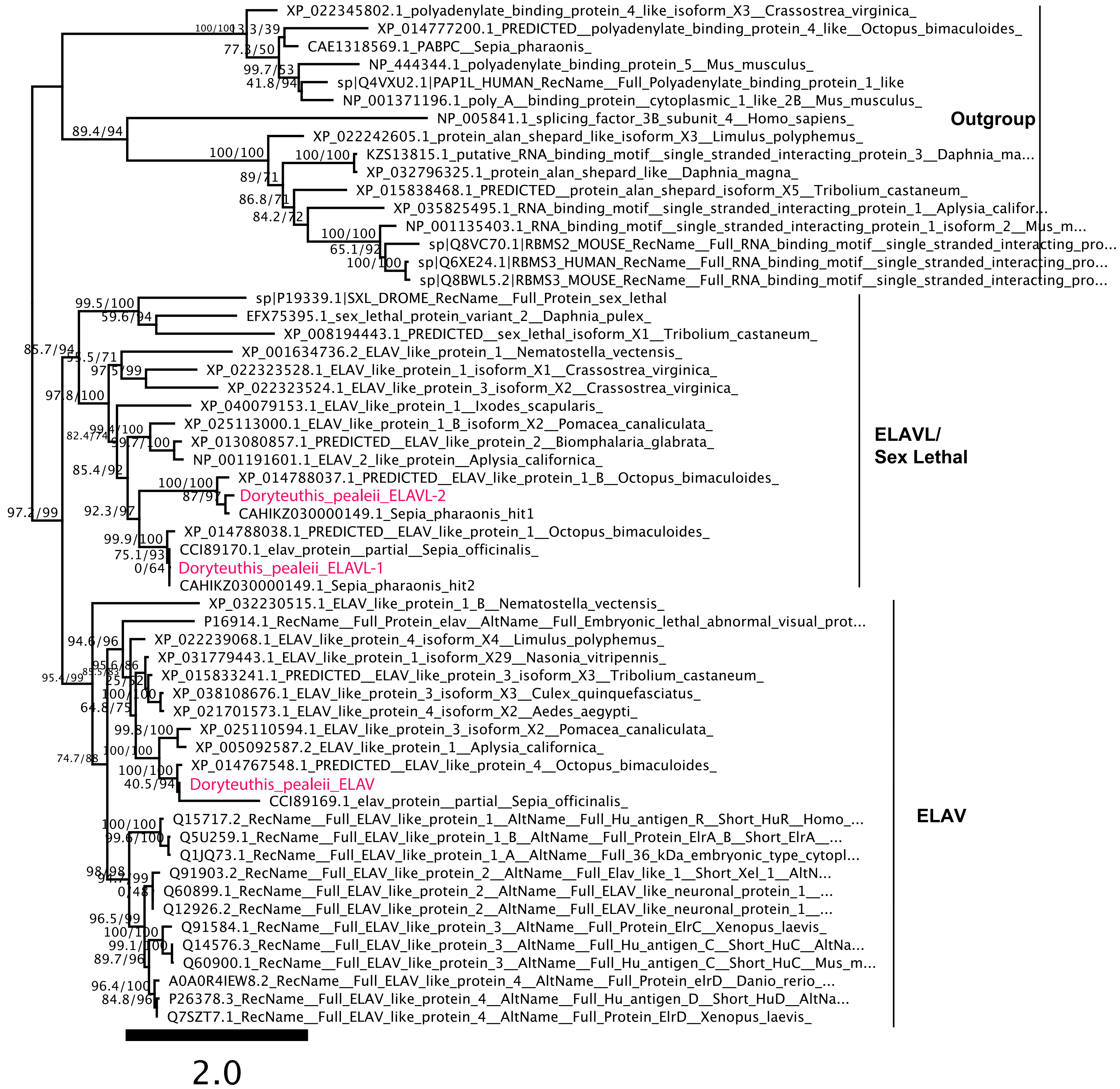

XP_022345802.1_polyadenylate_binding_protein_4_like_isoform_X3__Crassostrea_virginica_
XP_014777200.1_PREDICTED__polyadenylate_binding_protein_4_like__Octopus_bimaculoides_
CAE1318569.1_PABPC__Sepia_pharaonis_
NP_444344.1_polyadenylate_binding_protein_5__Mus_musculus_
sp|Q4VXU2.1|PAP1L_HUMAN_RecName__Full_Polyadenylate_binding_protein_1_like
NP_001371196.1_poly_A__binding_protein__cytoplasmic_1_like_2B__Mus_musculus_
NP_005841.1_splicing_factor_3B_subunit_4__Homo_sapiens_
XP_022242605.1_protein_alan_shepard_like_isoform_X3__Limulus_polyphemus_
KZS13815.1_putative_RNA_binding_motif__single_stranded_interacting_protein_3__Daphnia_ma...
XP_032796325.1_protein_alan_shepard_like__Daphnia_magna_
XP_015838468.1_PREDICTED__protein_alan_shepard_isoform_X5__Tribolium_castaneum_
XP_035825495.1_RNA_binding_motif__single_stranded_interacting_protein_1__Aplysia_califor...
NP_001135403.1_RNA_binding_motif__single_stranded_interacting_protein_1_isoform_2__Mus_m...
sp|Q8VC70.1|RBMS2_MOUSE_RecName__Full_RNA_binding_motif__single_stranded_interacting_pro...
sp|Q6XE24.1|RBMS3_HUMAN_RecName__Full_RNA_binding_motif__single_stranded_interacting_pro...
sp|Q8BWL5.2|RBMS3_MOUSE_RecName__Full_RNA_binding_motif__single_stranded_interacting_pro...
Outgroup
sp|P19339.1|SXL_DROME_RecName__Full_Protein_sex_lethal
EFX75395.1_sex_lethal_protein_variant_2__Daphnia_pulex_
XP_008194443.1_PREDICTED__sex_lethal_isoform_X1__Tribolium_castaneum_
XP_001634736.2_ELAV_like_protein_1__Nematostella_vectensis_
XP_022323528.1_ELAV_like_protein_1_isoform_X1__Crassostrea_virginica_
XP_022323524.1_ELAV_like_protein_3_isoform_X2__Crassostrea_virginica_
XP_040079153.1_ELAV_like_protein_1__Ixodes_scapularis_
XP_025113000.1_ELAV_like_protein_1_B_isoform_X2__Pomacea_canaliculata_
XP_013080857.1_PREDICTED__ELAV_like_protein_2__Biomphalaria_glabrata_
NP_001191601.1_ELAV_2_like_protein__Aplysia_californica_
XP_014788037.1_PREDICTED__ELAV_like_protein_1_B__Octopus_bimaculoides_
Doryteuthis_pealeii_ELAVL-2
CAHIKZ030000149.1_Sepia_pharaonis_hit1
XP_014788038.1_PREDICTED__ELAV_like_protein_1__Octopus_bimaculoides_
CCI89170.1_elav_protein__partial__Sepia_officinalis_
Doryteuthis_pealeii_ELAVL-1
CAHIKZ030000149.1_Sepia_pharaonis_hit2
ELAVL/
Sex Lethal
XP_032230515.1_ELAV_like_protein_1_B__Nematostella_vectensis_
P16914.1_RecName__Full_Protein_elav__AltName__Full_Embryonic_lethal_abnormal_visual_prot...
XP_022239068.1_ELAV_like_protein_4_isoform_X4__Limulus_polyphemus_
XP_031779443.1_ELAV_like_protein_1_isoform_X29__Nasonia_vitripennis_
XP_015833241.1_PREDICTED__ELAV_like_protein_3_isoform_X3__Tribolium_castaneum_
XP_038108676.1_ELAV_like_protein_3_isoform_X3__Culex_quinquefasciatus_
XP_021701573.1_ELAV_like_protein_4_isoform_X2__Aedes_aegypti_
XP_025110594.1_ELAV_like_protein_3_isoform_X2__Pomacea_canaliculata_
XP_005092587.2_ELAV_like_protein_1__Aplysia_californica_
XP_014767548.1_PREDICTED__ELAV_like_protein_4__Octopus_bimaculoides_
Doryteuthis_pealeii_ELAV
CCI89169.1_elav_protein__partial__Sepia_officinalis_
Q15717.2_RecName__Full_ELAV_like_protein_1__AltName__Full_Hu_antigen_R__Short_HuR__Homo_...
Q5U259.1_RecName__Full_ELAV_like_protein_1_B__AltName__Full_Protein_ElrA_B__Short_ElrA__...
Q1JQ73.1_RecName__Full_ELAV_like_protein_1_A__AltName__Full_36_kDa_embryonic_type_cytopl...
Q91903.2_RecName__Full_ELAV_like_protein_2__AltName__Full_Elav_like_1__Short_Xel_1__AltN...
Q60899.1_RecName__Full_ELAV_like_protein_2__AltName__Full_ELAV_like_neuronal_protein_1__...
Q12926.2_RecName__Full_ELAV_like_protein_2__AltName__Full_ELAV_like_neuronal_protein_1__...
Q91584.1_RecName__Full_ELAV_like_protein_3__AltName__Full_Protein_ElrC__Xenopus_laevis_
Q14576.3_RecName__Full_ELAV_like_protein_3__AltName__Full_Hu_antigen_C__Short_HuC__AltNa...
Q60900.1_RecName__Full_ELAV_like_protein_3__AltName__Full_Hu_antigen_C__Short_HuC__Mus_m...
A0A0R4IEW8.2_RecName__Full_ELAV_like_protein_4__AltName__Full_Protein_elrD__Danio_rerio_...
P26378.3_RecName__Full_ELAV_like_protein_4__AltName__Full_Hu_antigen_D__Short_HuD__AltNa...
Q7SZT7.1_RecName__Full_ELAV_like_protein_4__AltName__Full_Protein_ElrD__Xenopus_laevis_
ELAV
2.0

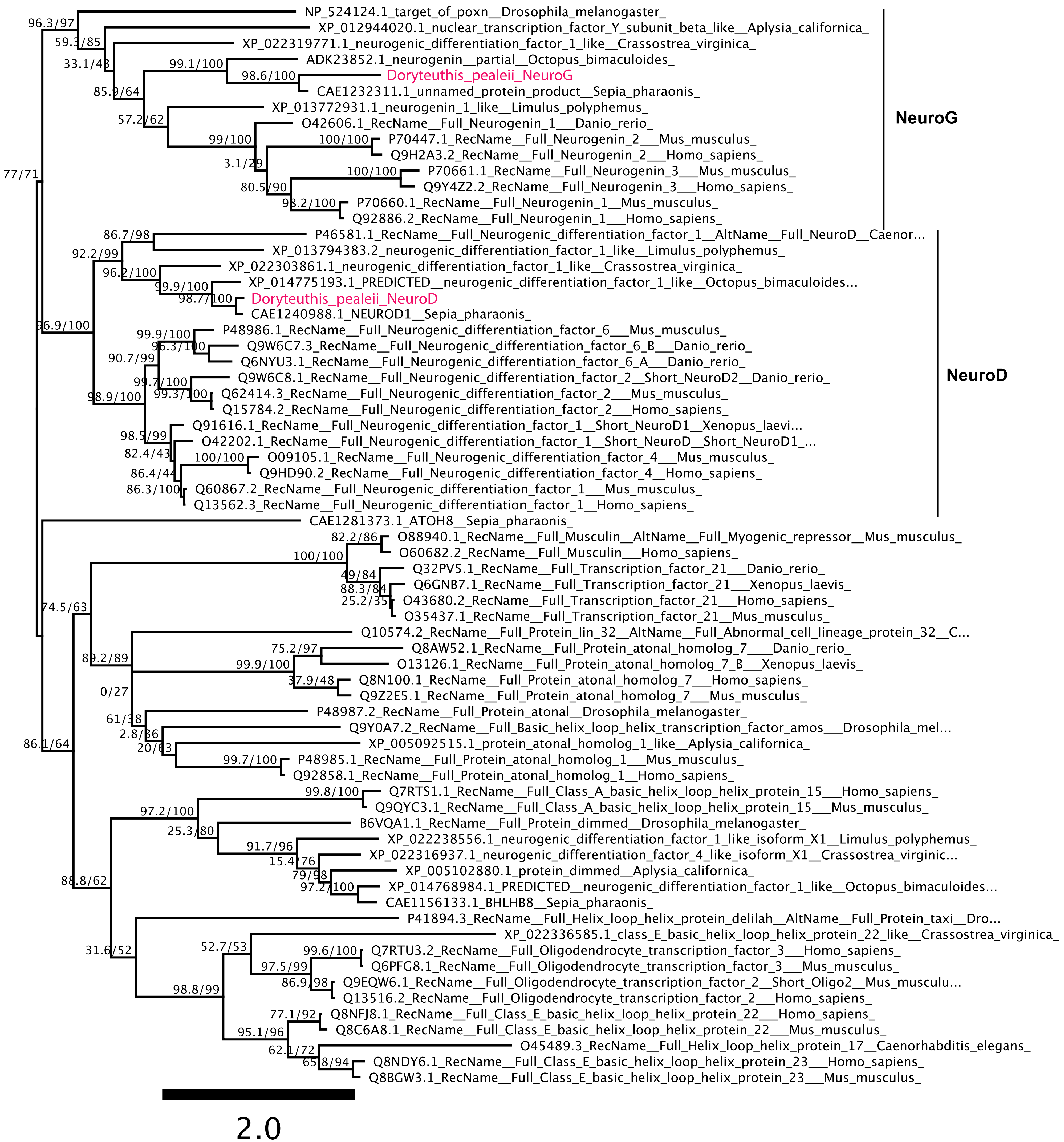
NP_524124.1_target_of_poxn__Drosophila_melanogaster_
XP_012944020.1_nuclear_transcription_factor_Y_subunit_beta_like__Aplysia_californica_
XP_022319771.1_neurogenic_differentiation_factor_1_like__Crassostrea_virginica_
ADK23852.1_neurogenin__partial__Octopus_bimaculoides_
Doryteuthis_pealeii_NeuroG
CAE1232311.1_unnamed_protein_product__Sepia_pharaonis_
XP_013772931.1_neurogenin_1_like__Limulus_polyphemus_
O42606.1_RecName__Full_Neurogenin_1___Danio_rerio_
P70447.1_RecName__Full_Neurogenin_2___Mus_musculus_
Q9H2A3.2_RecName__Full_Neurogenin_2___Homo_sapiens_
P70661.1_RecName__Full_Neurogenin_3___Mus_musculus_
Q9Y4Z2.2_RecName__Full_Neurogenin_3___Homo_sapiens_
P70660.1_RecName__Full_Neurogenin_1__Mus_musculus_
Q92886.2_RecName__Full_Neurogenin_1___Homo_sapiens_
NeuroG
P46581.1_RecName__Full_Neurogenic_differentiation_factor_1__AltName__Full_NeuroD__Caenor...
XP_013794383.2_neurogenic_differentiation_factor_1_like__Limulus_polyphemus_
XP_022303861.1_neurogenic_differentiation_factor_1_like__Crassostrea_virginica_
XP_014775193.1_PREDICTED__neurogenic_differentiation_factor_1_like__Octopus_bimaculoides...
Doryteuthis_pealeii_NeuroD
CAE1240988.1_NEUROD1__Sepia_pharaonis_
P48986.1_RecName__Full_Neurogenic_differentiation_factor_6___Mus_musculus_
Q9W6C7.3_RecName__Full_Neurogenic_differentiation_factor_6_B___Danio_rerio_
Q6NYU3.1_RecName__Full_Neurogenic_differentiation_factor_6_A___Danio_rerio_
Q9W6C8.1_RecName__Full_Neurogenic_differentiation_factor_2__Short_NeuroD2__Danio_rerio_
Q62414.3_RecName__Full_Neurogenic_differentiation_factor_2___Mus_musculus_
Q15784.2_RecName__Full_Neurogenic_differentiation_factor_2___Homo_sapiens_
Q91616.1_RecName__Full_Neurogenic_differentiation_factor_1__Short_NeuroD1__Xenopus_laevi...
O42202.1_RecName__Full_Neurogenic_differentiation_factor_1__Short_NeuroD__Short_NeuroD1_...
O09105.1_RecName__Full_Neurogenic_differentiation_factor_4___Mus_musculus_
Q9HD90.2_RecName__Full_Neurogenic_differentiation_factor_4__Homo_sapiens_
Q60867.2_RecName__Full_Neurogenic_differentiation_factor_1___Mus_musculus_
Q13562.3_RecName__Full_Neurogenic_differentiation_factor_1__Homo_sapiens_
NeuroD
CAE1281373.1_ATOH8__Sepia_pharaonis_
O88940.1_RecName__Full_Musculin__AltName__Full_Myogenic_repressor__Mus_musculus_
O60682.2_RecName__Full_Musculin___Homo_sapiens_
Q32PV5.1_RecName__Full_Transcription_factor_21___Danio_rerio_
Q6GNB7.1_RecName__Full_Transcription_factor_21___Xenopus_laevis_
O43680.2_RecName__Full_Transcription_factor_21___Homo_sapiens_
O35437.1_RecName__Full_Transcription_factor_21__Mus_musculus_
Q10574.2_RecName__Full_Protein_lin_32__AltName__Full_Abnormal_cell_lineage_protein_32__C...
Q8AW52.1_RecName__Full_Protein_atonal_homolog_7____Danio_rerio_
O13126.1_RecName__Full_Protein_atonal_homolog_7_B___Xenopus_laevis_
Q8N100.1_RecName__Full_Protein_atonal_homolog_7___Homo_sapiens_
Q9Z2E5.1_RecName__Full_Protein_atonal_homolog_7___Mus_musculus_
P48987.2_RecName__Full_Protein_atonal__Drosophila_melanogaster_
Q9Y0A7.2_RecName__Full_Basic_helix_loop_helix_transcription_factor_amos___Drosophila_mel...
XP_005092515.1_protein_atonal_homolog_1_like__Aplysia_californica_
P48985.1_RecName__Full_Protein_atonal_homolog_1___Mus_musculus_
Q92858.1_RecName__Full_Protein_atonal_homolog_1__Homo_sapiens_
Q7RTS1.1_RecName__Full_Class_A_basic_helix_loop_helix_protein_15___Homo_sapiens_
Q9QYC3.1_RecName__Full_Class_A_basic_helix_loop_helix_protein_15___Mus_musculus_
B6VQA1.1_RecName__Full_Protein_dimmed__Drosophila_melanogaster_
XP_022238556.1_neurogenic_differentiation_factor_1_like_isoform_X1__Limulus_polyphemus_
XP_022316937.1_neurogenic_differentiation_factor_4_like_isoform_X1__Crassostrea_virginic...
XP_005102880.1_protein_dimmed__Aplysia_californica_
XP_014768984.1_PREDICTED__neurogenic_differentiation_factor_1_like__Octopus_bimaculoides...
CAE1156133.1_BHLHB8__Sepia_pharaonis_
P41894.3_RecName__Full_Helix_loop_helix_protein_delilah__AltName__Full_Protein_taxi__Dro...
XP_022336585.1_class_E_basic_helix_loop_helix_protein_22_like__Crassostrea_virginica_
Q7RTU3.2_RecName__Full_Oligodendrocyte_transcription_factor_3___Homo_sapiens_
Q6PFG8.1_RecName__Full_Oligodendrocyte_transcription_factor_3___Mus_musculus_
Q9EQW6.1_RecName__Full_Oligodendrocyte_transcription_factor_2__Short_Oligo2__Mus_musculu...
Q13516.2_RecName__Full_Oligodendrocyte_transcription_factor_2___Homo_sapiens_
Q8NFJ8.1_RecName__Full_Class_E_basic_helix_loop_helix_protein_22___Homo_sapiens_
Q8C6A8.1_RecName__Full_Class_E_basic_helix_loop_helix_protein_22___Mus_musculus_
O45489.3_RecName__Full_Helix_loop_helix_protein_17__Caenorhabditis_elegans_
Q8NDY6.1_RecName__Full_Class_E_basic_helix_loop_helix_protein_23___Homo_sapiens_
Q8BGW3.1_RecName__Full_Class_E_basic_helix_loop_helix_protein_23___Mus_musculus_
2.0

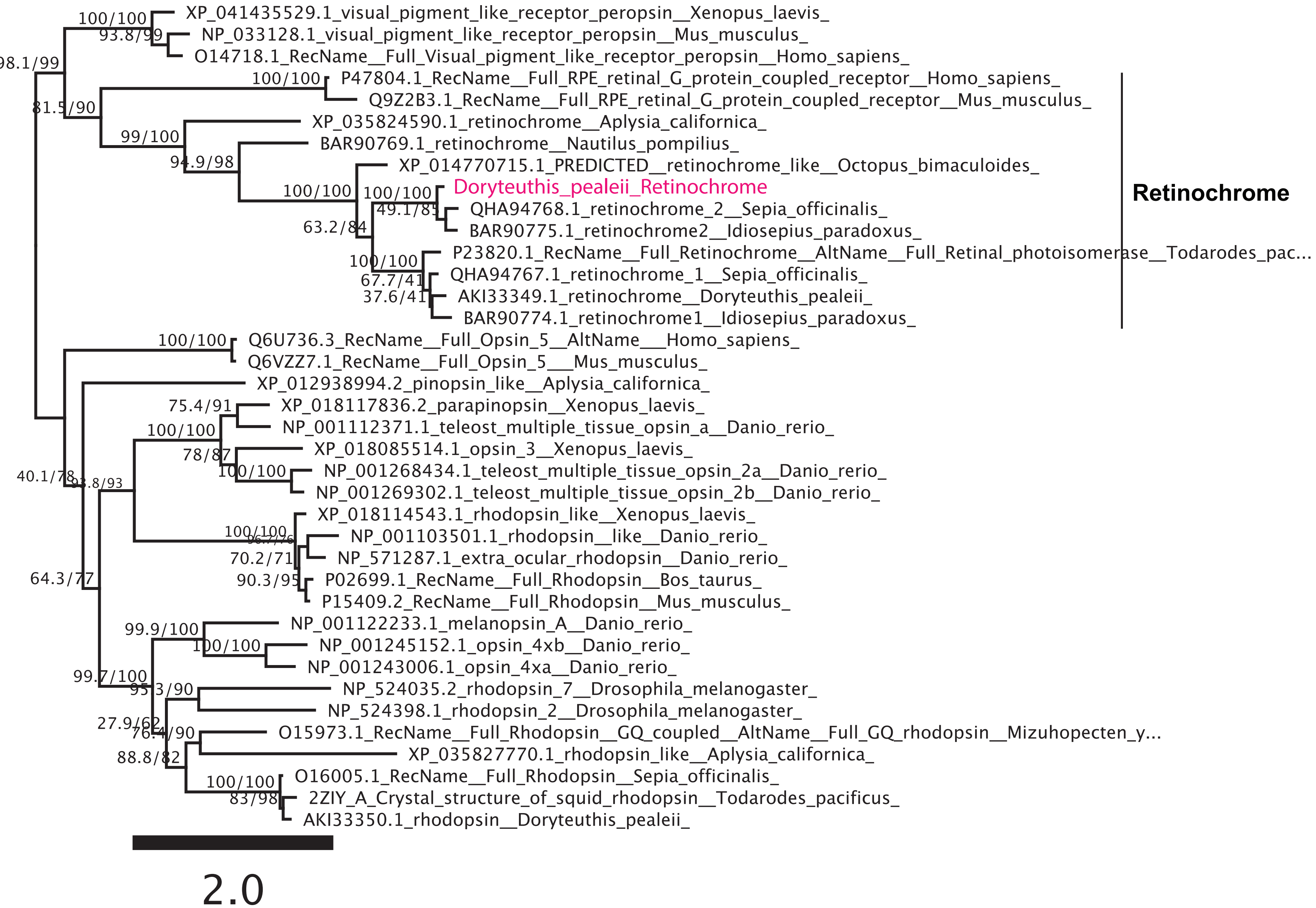

XP_041435529.1_visual_pigment_like_receptor_peropsin__Xenopus_laevis_
NP_033128.1_visual_pigment_like_receptor_peropsin__Mus_musculus_
O14718.1_RecName__Full_Visual_pigment_like_receptor_peropsin__Homo_sapiens_
P47804.1_RecName__Full_RPE_retinal_G_protein_coupled_receptor__Homo_sapiens_
Q9Z2B3.1_RecName__Full_RPE_retinal_G_protein_coupled_receptor__Mus_musculus_
XP_035824590.1_retinochrome__Aplysia_californica_
BAR90769.1_retinochrome__Nautilus_pompilius_
XP_014770715.1_PREDICTED__retinochrome_like__Octopus_bimaculoides_
Doryteuthis_pealeii_Retinochrome
QHA94768.1_retinochrome_2__Sepia_officinalis_
BAR90775.1_retinochrome2__Idiosepius_paradoxus_
P23820.1_RecName__Full_Retinochrome__AltName__Full_Retinal_photoisomerase__Todarodes_pac...
QHA94767.1_retinochrome_1__Sepia_officinalis_
AKI33349.1_retinochrome__Doryteuthis_pealeii_
BAR90774.1_retinochrome1__Idiosepius_paradoxus_
Retinochrome
Q6U736.3_RecName__Full_Opsin_5__AltName___Homo_sapiens_
Q6VZZ7.1_RecName__Full_Opsin_5___Mus_musculus_
XP_012938994.2_pinopsin_like__Aplysia_californica_
XP_018117836.2_parapinopsin__Xenopus_laevis_
NP_001112371.1_teleost_multiple_tissue_opsin_a__Danio_rerio_
XP_018085514.1_opsin_3__Xenopus_laevis_
NP_001268434.1_teleost_multiple_tissue_opsin_2a__Danio_rerio_
NP_001269302.1_teleost_multiple_tissue_opsin_2b__Danio_rerio_
XP_018114543.1_rhodopsin_like__Xenopus_laevis_
NP_001103501.1_rhodopsin__like__Danio_rerio_
NP_571287.1_extra_ocular_rhodopsin__Danio_rerio_
P02699.1_RecName__Full_Rhodopsin__Bos_taurus_
P15409.2_RecName__Full_Rhodopsin__Mus_musculus_
NP_001122233.1_melanopsin_A__Danio_rerio_
NP_001245152.1_opsin_4xb__Danio_rerio_
NP_001243006.1_opsin_4xa__Danio_rerio_
NP_524035.2_rhodopsin_7__Drosophila_melanogaster_
NP_524398.1_rhodopsin_2__Drosophila_melanogaster_
O15973.1_RecName__Full_Rhodopsin__GQ_coupled__AltName__Full_GQ_rhodopsin__Mizuhopecten_y...
XP_035827770.1_rhodopsin_like__Aplysia_californica_
O16005.1_RecName__Full_Rhodopsin__Sepia_officinalis_
2ZIY_A_Crystal_structure_of_squid_rhodopsin__Todarodes_pacificus_
AKI33350.1_rhodopsin__Doryteuthis_pealeii_
100/100
93.8/99
98.1/99
81.5/90
99/100
94.9/98
100/100
63.2/84
49.1/85
67.7/41
37.6/41
40.1/78
64.3/77
75.4/91
78/87
70.2/71
90.3/95
99.9/100
99.7/100
95.3/90
27.9/62
76.4/90
88.8/82
83/98
2.0

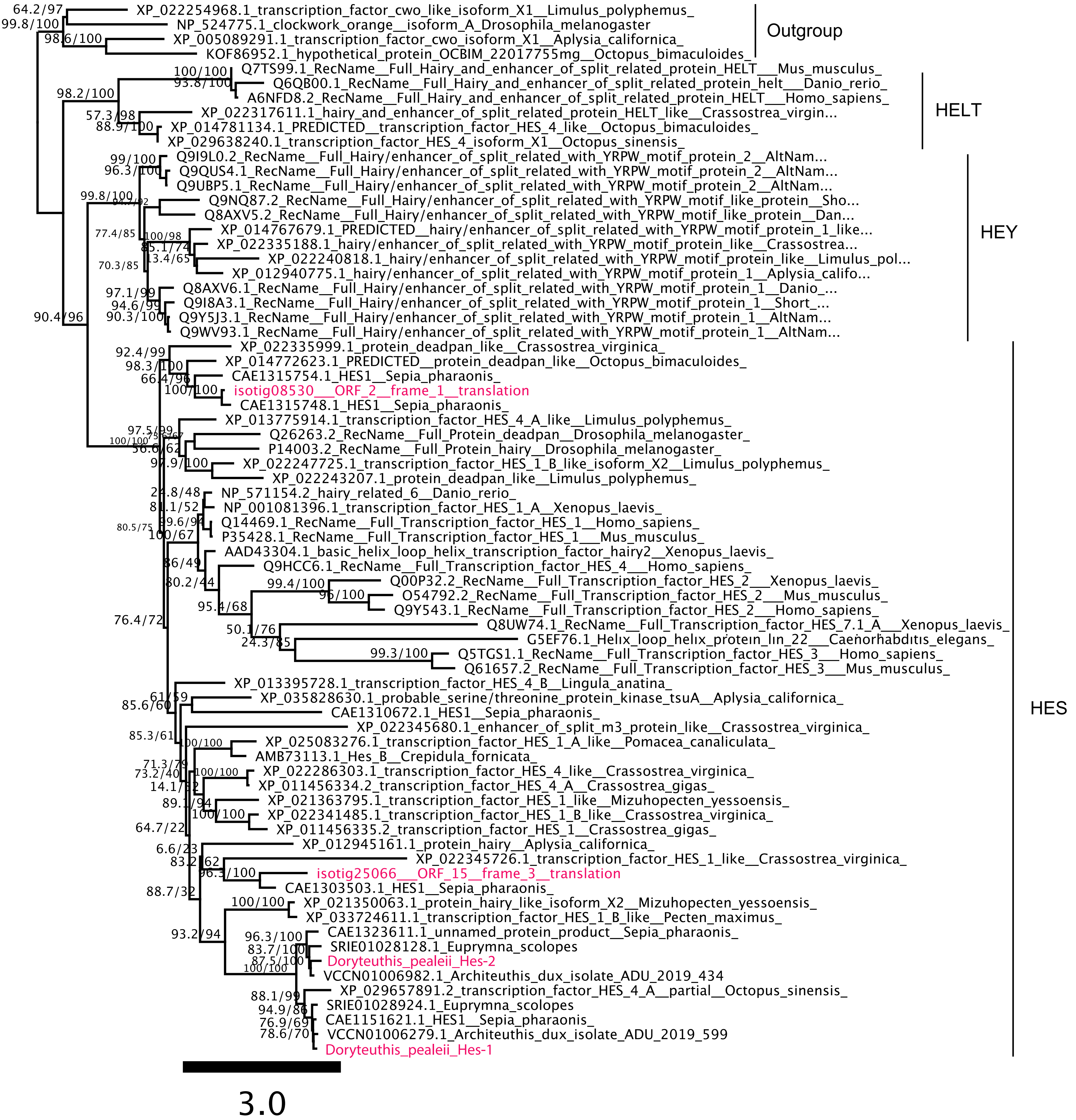

XP_022254968.1_transcription_factor_cwo_like_isoform_X1__Limulus_polyphemus_
NP_524775.1_clockwork_orange__isoform_A_Drosophila_melanogaster
XP_005089291.1_transcription_factor_cwo_isoform_X1__Aplysia_californica_
KOF86952.1_hypothetical_protein_OCBIM_22017755mg__Octopus_bimaculoides_
Outgroup
Q7TS99.1_RecName__Full_Hairy_and_enhancer_of_split_related_protein_HELT___Mus_musculus_
Q6QB00.1_RecName__Full_Hairy_and_enhancer_of_split_related_protein_helt___Danio_rerio_
A6NFD8.2_RecName__Full_Hairy_and_enhancer_of_split_related_protein_HELT___Homo_sapiens_
XP_022317611.1_hairy_and_enhancer_of_split_related_protein_HELT_like__Crassostrea_virgin...
XP_014781134.1_PREDICTED__transcription_factor_HES_4_like__Octopus_bimaculoides_
XP_029638240.1_transcription_factor_HES_4_isoform_X1__Octopus_sinensis_
HELT
Q9I9L0.2_RecName__Full_Hairy/enhancer_of_split_related_with_YRPW_motif_protein_2__AltNam...
Q9QUS4.1_RecName__Full_Hairy/enhancer_of_split_related_with_YRPW_motif_protein_2__AltNam...
Q9UBP5.1_RecName__Full_Hairy/enhancer_of_split_related_with_YRPW_motif_protein_2__AltNam...
Q9NQ87.2_RecName__Full_Hairy/enhancer_of_split_related_with_YRPW_motif_like_protein__Sho...
Q8AXV5.2_RecName__Full_Hairy/enhancer_of_split_related_with_YRPW_motif_like_protein__Dan...
XP_014767679.1_PREDICTED__hairy/enhancer_of_split_related_with_YRPW_motif_protein_1_like...
XP_022335188.1_hairy/enhancer_of_split_related_with_YRPW_motif_protein_like__Crassostrea...
XP_022240818.1_hairy/enhancer_of_split_related_with_YRPW_motif_protein_like__Limulus_pol...
XP_012940775.1_hairy/enhancer_of_split_related_with_YRPW_motif_protein_1__Aplysia_califo...
Q8AXV6.1_RecName__Full_Hairy/enhancer_of_split_related_with_YRPW_motif_protein_1__Danio_...
Q9I8A3.1_RecName__Full_Hairy/enhancer_of_split_related_with_YRPW_motif_protein_1__Short_...
Q9Y5J3.1_RecName__Full_Hairy/enhancer_of_split_related_with_YRPW_motif_protein_1__AltNam...
Q9WV93.1_RecName__Full_Hairy/enhancer_of_split_related_with_YRPW_motif_protein_1__AltNam...
HEY
XP_022335999.1_protein_deadpan_like__Crassostrea_virginica_
XP_014772623.1_PREDICTED__protein_deadpan_like__Octopus_bimaculoides_
CAE1315754.1_HES1__Sepia_pharaonis_
isotig08530___ORF_2__frame_1__translation
CAE1315748.1_HES1__Sepia_pharaonis_
XP_013775914.1_transcription_factor_HES_4_A_like__Limulus_polyphemus_
Q26263.2_RecName__Full_Protein_deadpan__Drosophila_melanogaster_
P14003.2_RecName__Full_Protein_hairy__Drosophila_melanogaster_
XP_022247725.1_transcription_factor_HES_1_B_like_isoform_X2__Limulus_polyphemus_
XP_022243207.1_protein_deadpan_like__Limulus_polyphemus_
NP_571154.2_hairy_related_6__Danio_rerio_
NP_001081396.1_transcription_factor_HES_1_A__Xenopus_laevis_
Q14469.1_RecName__Full_Transcription_factor_HES_1__Homo_sapiens_
P35428.1_RecName__Full_Transcription_factor_HES_1___Mus_musculus_
AAD43304.1_basic_helix_loop_helix_transcription_factor_hairy2__Xenopus_laevis_
Q9HCC6.1_RecName__Full_Transcription_factor_HES_4___Homo_sapiens_
Q00P32.2_RecName__Full_Transcription_factor_HES_2__Xenopus_laevis_
O54792.2_RecName__Full_Transcription_factor_HES_2__Mus_musculus_
Q9Y543.1_RecName__Full_Transcription_factor_HES_2__Homo_sapiens_
Q8UW74.1_RecName__Full_Transcription_factor_HES_7.1_A___Xenopus_laevis_
G5EF76.1_Helix_loop_helix_protein_lin_22___Caenorhabditis_elegans_
Q5TGS1.1_RecName__Full_Transcription_factor_HES_3___Homo_sapiens_
Q61657.2_RecName__Full_Transcription_factor_HES_3___Mus_musculus_
XP_013395728.1_transcription_factor_HES_4_B__Lingula_anatina_
XP_035828630.1_probable_serine/threonine_protein_kinase_tsuA__Aplysia_californica_
CAE1310672.1_HES1__Sepia_pharaonis_
XP_022345680.1_enhancer_of_split_m3_protein_like__Crassostrea_virginica_
XP_025083276.1_transcription_factor_HES_1_A_like__Pomacea_canaliculata_
AMB73113.1_Hes_B__Crepidula_fornicata_
XP_022286303.1_transcription_factor_HES_4_like__Crassostrea_virginica_
XP_011456334.2_transcription_factor_HES_4_A__Crassostrea_gigas_
XP_021363795.1_transcription_factor_HES_1_like__Mizuhopecten_yessoensis_
XP_022341485.1_transcription_factor_HES_1_B_like__Crassostrea_virginica_
XP_011456335.2_transcription_factor_HES_1__Crassostrea_gigas_
XP_012945161.1_protein_hairy__Aplysia_californica_
XP_022345726.1_transcription_factor_HES_1_like__Crassostrea_virginica_
isotig25066___ORF_15__frame_3__translation
CAE1303503.1_HES1__Sepia_pharaonis_
XP_021350063.1_protein_hairy_like_isoform_X2__Mizuhopecten_yessoensis_
XP_033724611.1_transcription_factor_HES_1_B_like__Pecten_maximus_
CAE1323611.1_unnamed_protein_product__Sepia_pharaonis_
SRIE01028128.1_Euprymna_scolopes
Doryteuthis_pealeii_Hes-2
VCCN01006982.1_Architeuthis_dux_isolate_ADU_2019_434
XP_029657891.2_transcription_factor_HES_4_A__partial__Octopus_sinensis_
SRIE01028924.1_Euprymna_scolopes
CAE1151621.1_HES1__Sepia_pharaonis_
VCCN01006279.1_Architeuthis_dux_isolate_ADU_2019_599
Doryteuthis_pealeii_Hes-1
HES
3.0

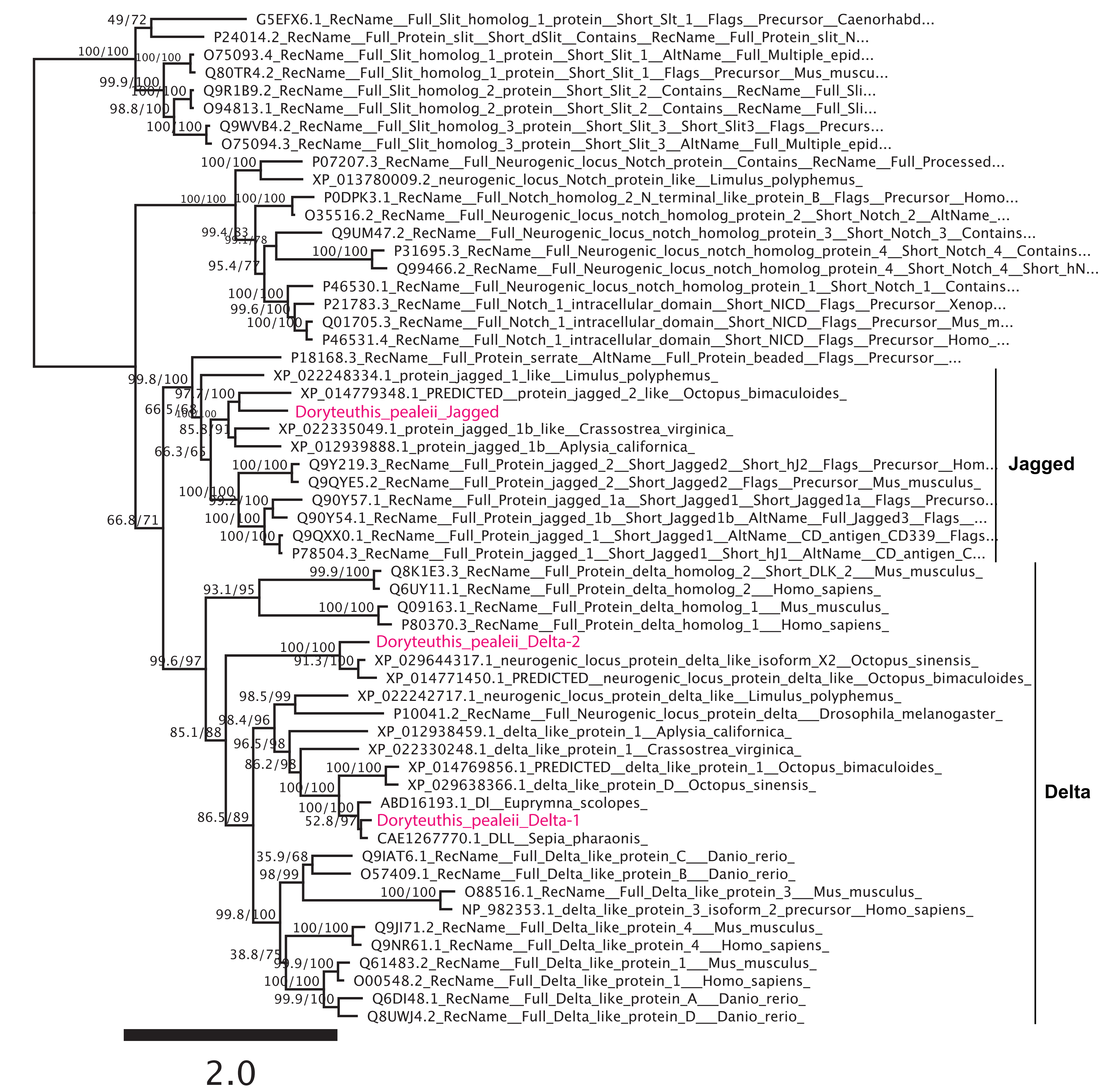
G5EFX6.1_RecName__Full_Slit_homolog_1_protein__Short_Slt_1__Flags__Precursor__Caenorhabd...
P24014.2_RecName__Full_Protein_slit__Short_dSlit__Contains__RecName__Full_Protein_slit_N...
O75093.4_RecName__Full_Slit_homolog_1_protein__Short_Slit_1__AltName__Full_Multiple_epid...
Q80TR4.2_RecName__Full_Slit_homolog_1_protein__Short_Slit_1__Flags__Precursor__Mus_muscu...
Q9R1B9.2_RecName__Full_Slit_homolog_2_protein__Short_Slit_2__Contains__RecName__Full_Sli...
O94813.1_RecName__Full_Slit_homolog_2_protein__Short_Slit_2__Contains__RecName__Full_Sli...
Q9WVB4.2_RecName__Full_Slit_homolog_3_protein__Short_Slit_3__Short_Slit3__Flags__Precurs...
O75094.3_RecName__Full_Slit_homolog_3_protein__Short_Slit_3__AltName__Full_Multiple_epid...
P07207.3_RecName__Full_Neurogenic_locus_Notch_protein__Contains__RecName__Full_Processed...
XP_013780009.2_neurogenic_locus_Notch_protein_like__Limulus_polyphemus_
P0DPK3.1_RecName__Full_Notch_homolog_2_N_terminal_like_protein_B__Flags__Precursor__Homo...
O35516.2_RecName__Full_Neurogenic_locus_notch_homolog_protein_2__Short_Notch_2__AltName_...
Q9UM47.2_RecName__Full_Neurogenic_locus_notch_homolog_protein_3__Short_Notch_3__Contains...
P31695.3_RecName__Full_Neurogenic_locus_notch_homolog_protein_4__Short_Notch_4__Contains...
Q99466.2_RecName__Full_Neurogenic_locus_notch_homolog_protein_4__Short_Notch_4__Short_hN...
P46530.1_RecName__Full_Neurogenic_locus_notch_homolog_protein_1__Short_Notch_1__Contains...
P21783.3_RecName__Full_Notch_1_intracellular_domain__Short_NICD__Flags__Precursor__Xenop...
Q01705.3_RecName__Full_Notch_1_intracellular_domain__Short_NICD__Flags__Precursor__Mus_m...
P46531.4_RecName__Full_Notch_1_intracellular_domain__Short_NICD__Flags__Precursor__Homo_...
P18168.3_RecName__Full_Protein_serrate__AltName__Full_Protein_beaded__Flags__Precursor__...
XP_022248334.1_protein_jagged_1_like__Limulus_polyphemus_
XP_014779348.1_PREDICTED__protein_jagged_2_like__Octopus_bimaculoides_
Doryteuthis_pealeii_Jagged
XP_022335049.1_protein_jagged_1b_like__Crassostrea_virginica_
XP_012939888.1_protein_jagged_1b__Aplysia_californica_
Q9Y219.3_RecName__Full_Protein_jagged_2__Short_Jagged2__Short_hJ2__Flags__Precursor__Hom...
Q9QYE5.2_RecName__Full_Protein_jagged_2__Short_Jagged2__Flags__Precursor__Mus_musculus_
Q90Y57.1_RecName__Full_Protein_jagged_1a__Short_Jagged1__Short_Jagged1a__Flags__Precurso...
Q90Y54.1_RecName__Full_Protein_jagged_1b__Short_Jagged1b__AltName__Full_Jagged3__Flags__...
Q9QXX0.1_RecName__Full_Protein_jagged_1__Short_Jagged1__AltName__CD_antigen_CD339__Flags...
P78504.3_RecName__Full_Protein_jagged_1__Short_Jagged1__Short_hJ1__AltName__CD_antigen_C...
Jagged
Q8K1E3.3_RecName__Full_Protein_delta_homolog_2__Short_DLK_2___Mus_musculus_
Q6UY11.1_RecName__Full_Protein_delta_homolog_2___Homo_sapiens_
Q09163.1_RecName__Full_Protein_delta_homolog_1___Mus_musculus_
P80370.3_RecName__Full_Protein_delta_homolog_1___Homo_sapiens_
Doryteuthis_pealeii_Delta-2
XP_029644317.1_neurogenic_locus_protein_delta_like_isoform_X2__Octopus_sinensis_
XP_014771450.1_PREDICTED__neurogenic_locus_protein_delta_like__Octopus_bimaculoides_
XP_022242717.1_neurogenic_locus_protein_delta_like__Limulus_polyphemus_
P10041.2_RecName__Full_Neurogenic_locus_protein_delta___Drosophila_melanogaster_
XP_012938459.1_delta_like_protein_1__Aplysia_californica_
XP_022330248.1_delta_like_protein_1__Crassostrea_virginica_
XP_014769856.1_PREDICTED__delta_like_protein_1__Octopus_bimaculoides_
XP_029638366.1_delta_like_protein_D__Octopus_sinensis_
ABD16193.1_Dl__Euprymna_scolopes_
Doryteuthis_pealeii_Delta-1
CAE1267770.1_DLL__Sepia_pharaonis_
Q9IAT6.1_RecName__Full_Delta_like_protein_C___Danio_rerio_
O57409.1_RecName__Full_Delta_like_protein_B___Danio_rerio_
O88516.1_RecName__Full_Delta_like_protein_3___Mus_musculus_
NP_982353.1_delta_like_protein_3_isoform_2_precursor__Homo_sapiens_
Q9JI71.2_RecName__Full_Delta_like_protein_4___Mus_musculus_
Q9NR61.1_RecName__Full_Delta_like_protein_4___Homo_sapiens_
Q61483.2_RecName__Full_Delta_like_protein_1___Mus_musculus_
O00548.2_RecName__Full_Delta_like_protein_1___Homo_sapiens_
Q6DI48.1_RecName__Full_Delta_like_protein_A___Danio_rerio_
Q8UWJ4.2_RecName__Full_Delta_like_protein_D___Danio_rerio_
Delta
2.0

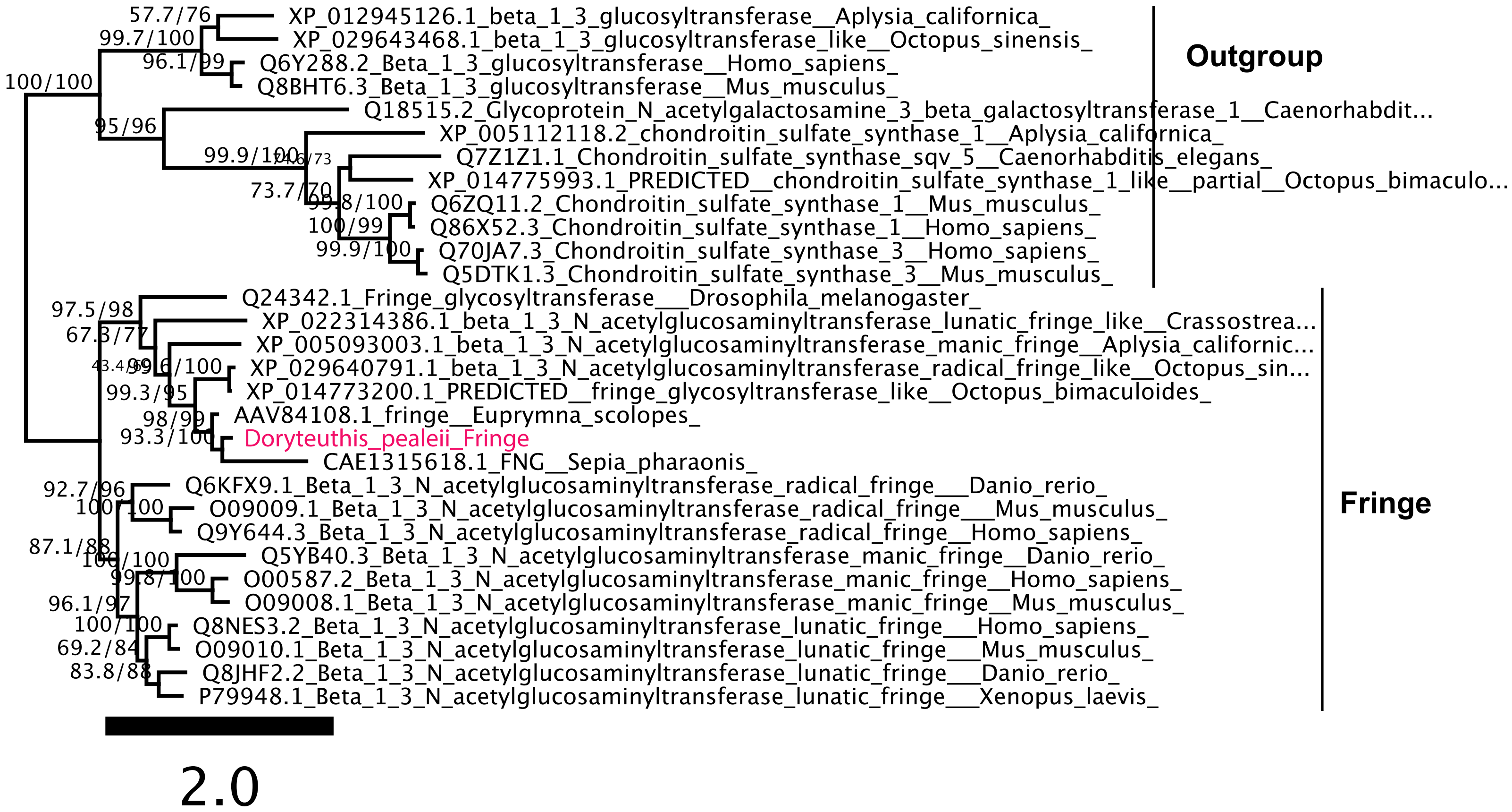

57.7/76
XP_012945126.1_beta_1_3_glucosyltransferase__Aplysia_californica_
99.7/100
XP_029643468.1_beta_1_3_glucosyltransferase_like__Octopus_sinensis_
96.1/99
Q6Y288.2_Beta_1_3_glucosyltransferase__Homo_sapiens_
100/100
Q8BHT6.3_Beta_1_3_glucosyltransferase__Mus_musculus_
Q18515.2_Glycoprotein_N_acetylgalactosamine_3_beta_galactosyltransferase_1__Caenorhabdit...
95/96
XP_005112118.2_chondroitin_sulfate_synthase_1__Aplysia_californica_
99.9/100
74.6/73
Q7Z1Z1.1_Chondroitin_sulfate_synthase_sqv_5__Caenorhabditis_elegans_
XP_014775993.1_PREDICTED__chondroitin_sulfate_synthase_1_like__partial__Octopus_bimaculo...
73.7/70
99.8/100
Q6ZQ11.2_Chondroitin_sulfate_synthase_1__Mus_musculus_
100/99
Q86X52.3_Chondroitin_sulfate_synthase_1__Homo_sapiens_
99.9/100
Q70JA7.3_Chondroitin_sulfate_synthase_3__Homo_sapiens_
Q5DTK1.3_Chondroitin_sulfate_synthase_3__Mus_musculus_
Outgroup
Q24342.1_Fringe_glycosyltransferase___Drosophila_melanogaster_
97.5/98
XP_022314386.1_beta_1_3_N_acetylglucosaminyltransferase_lunatic_fringe_like__Crassostrea...
67.3/77
XP_005093003.1_beta_1_3_N_acetylglucosaminyltransferase_manic_fringe__Aplysia_californic...
43.4/39
99.6/100
XP_029640791.1_beta_1_3_N_acetylglucosaminyltransferase_radical_fringe_like__Octopus_sin...
99.3/95
XP_014773200.1_PREDICTED__fringe_glycosyltransferase_like__Octopus_bimaculoides_
98/99
AAV84108.1_fringe__Euprymna_scolopes_
93.3/100
Doryteuthis_pealeii_Fringe
CAE1315618.1_FNG__Sepia_pharaonis_
92.7/96
Q6KFX9.1_Beta_1_3_N_acetylglucosaminyltransferase_radical_fringe___Danio_rerio_
100/100
O09009.1_Beta_1_3_N_acetylglucosaminyltransferase_radical_fringe___Mus_musculus_
Q9Y644.3_Beta_1_3_N_acetylglucosaminyltransferase_radical_fringe__Homo_sapiens_
87.1/88
100/100
Q5YB40.3_Beta_1_3_N_acetylglucosaminyltransferase_manic_fringe__Danio_rerio_
99.8/100
O00587.2_Beta_1_3_N_acetylglucosaminyltransferase_manic_fringe__Homo_sapiens_
96.1/97
O09008.1_Beta_1_3_N_acetylglucosaminyltransferase_manic_fringe__Mus_musculus_
100/100
Q8NES3.2_Beta_1_3_N_acetylglucosaminyltransferase_lunatic_fringe___Homo_sapiens_
69.2/84
O09010.1_Beta_1_3_N_acetylglucosaminyltransferase_lunatic_fringe___Mus_musculus_
83.8/88
Q8JHF2.2_Beta_1_3_N_acetylglucosaminyltransferase_lunatic_fringe___Danio_rerio_
P79948.1_Beta_1_3_N_acetylglucosaminyltransferase_lunatic_fringe___Xenopus_laevis_
Fringe
2.0

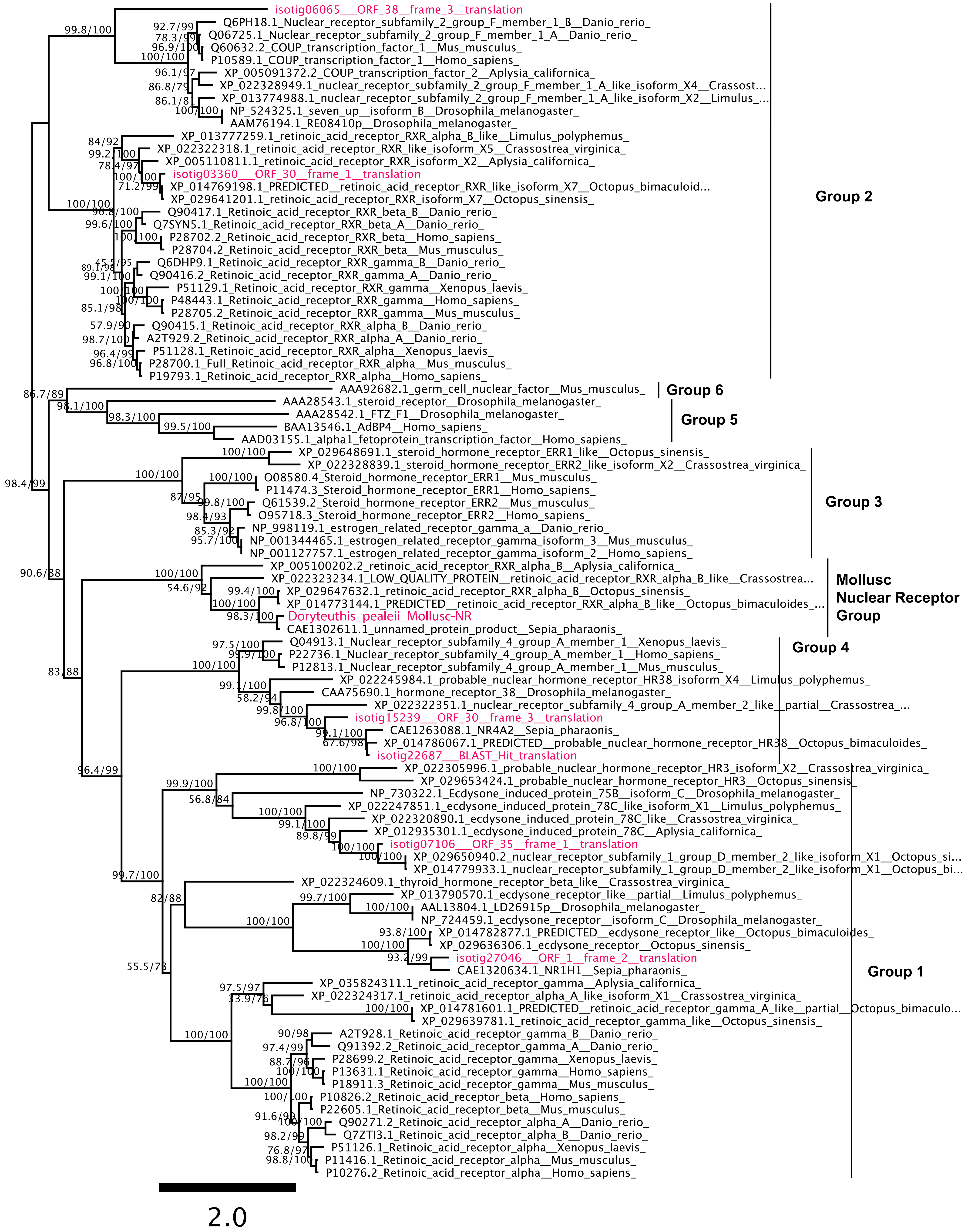
isotig06065___ORF_38__frame_3__translation
Q6PH18.1_Nuclear_receptor_subfamily_2_group_F_member_1_B__Danio_rerio_
Q06725.1_Nuclear_receptor_subfamily_2_group_F_member_1_A__Danio_rerio_
Q60632.2_COUP_transcription_factor_1__Mus_musculus_
P10589.1_COUP_transcription_factor_1__Homo_sapiens_
XP_005091372.2_COUP_transcription_factor_2__Aplysia_californica_
XP_022328949.1_nuclear_receptor_subfamily_2_group_F_member_1_A_like_isoform_X4__Crassost...
XP_013774988.1_nuclear_receptor_subfamily_2_group_F_member_1_A_like_isoform_X2__Limulus_...
NP_524325.1_seven_up__isoform_B__Drosophila_melanogaster_
AAM76194.1_RE08410p__Drosophila_melanogaster_
XP_013777259.1_retinoic_acid_receptor_RXR_alpha_B_like__Limulus_polyphemus_
XP_022322318.1_retinoic_acid_receptor_RXR_like_isoform_X5__Crassostrea_virginica_
XP_005110811.1_retinoic_acid_receptor_RXR_isoform_X2__Aplysia_californica_
isotig03360___ORF_30__frame_1__translation
XP_014769198.1_PREDICTED__retinoic_acid_receptor_RXR_like_isoform_X7__Octopus_bimaculoid...
XP_029641201.1_retinoic_acid_receptor_RXR_isoform_X7__Octopus_sinensis_
Q90417.1_Retinoic_acid_receptor_RXR_beta_B__Danio_rerio_
Q7SYN5.1_Retinoic_acid_receptor_RXR_beta_A__Danio_rerio_
P28702.2_Retinoic_acid_receptor_RXR_beta__Homo_sapiens_
P28704.2_Retinoic_acid_receptor_RXR_beta__Mus_musculus_
Q6DHP9.1_Retinoic_acid_receptor_RXR_gamma_B__Danio_rerio_
Q90416.2_Retinoic_acid_receptor_RXR_gamma_A__Danio_rerio_
P51129.1_Retinoic_acid_receptor_RXR_gamma__Xenopus_laevis_
P48443.1_Retinoic_acid_receptor_RXR_gamma__Homo_sapiens_
P28705.2_Retinoic_acid_receptor_RXR_gamma__Mus_musculus_
Q90415.1_Retinoic_acid_receptor_RXR_alpha_B__Danio_rerio_
A2T929.2_Retinoic_acid_receptor_RXR_alpha_A__Danio_rerio_
P51128.1_Retinoic_acid_receptor_RXR_alpha__Xenopus_laevis_
P28700.1_Full_Retinoic_acid_receptor_RXR_alpha__Mus_musculus_
P19793.1_Retinoic_acid_receptor_RXR_alpha__Homo_sapiens_
Group 2
AAA92682.1_germ_cell_nuclear_factor__Mus_musculus_
Group 6
AAA28543.1_steroid_receptor__Drosophila_melanogaster_
AAA28542.1_FTZ_F1__Drosophila_melanogaster_
BAA13546.1_AdBP4__Homo_sapiens_
AAD03155.1_alpha1_fetoprotein_transcription_factor__Homo_sapiens_
Group 5
XP_029648691.1_steroid_hormone_receptor_ERR1_like__Octopus_sinensis_
XP_022328839.1_steroid_hormone_receptor_ERR2_like_isoform_X2__Crassostrea_virginica_
O08580.4_Steroid_hormone_receptor_ERR1__Mus_musculus_
P11474.3_Steroid_hormone_receptor_ERR1__Homo_sapiens_
Q61539.2_Steroid_hormone_receptor_ERR2__Mus_musculus_
O95718.3_Steroid_hormone_receptor_ERR2__Homo_sapiens_
NP_998119.1_estrogen_related_receptor_gamma_a__Danio_rerio_
NP_001344465.1_estrogen_related_receptor_gamma_isoform_3__Mus_musculus_
NP_001127757.1_estrogen_related_receptor_gamma_isoform_2__Homo_sapiens_
Group 3
XP_005100202.2_retinoic_acid_receptor_RXR_alpha_B__Aplysia_californica_
XP_022323234.1_LOW_QUALITY_PROTEIN__retinoic_acid_receptor_RXR_alpha_B_like__Crassostrea...
XP_029647632.1_retinoic_acid_receptor_RXR_alpha_B__Octopus_sinensis_
XP_014773144.1_PREDICTED__retinoic_acid_receptor_RXR_alpha_B_like__Octopus_bimaculoides_...
Doryteuthis_pealeii_Mollusc-NR
CAE1302611.1_unnamed_protein_product__Sepia_pharaonis_
Mollusc Nuclear Receptor Group
Q04913.1_Nuclear_receptor_subfamily_4_group_A_member_1__Xenopus_laevis_
P22736.1_Nuclear_receptor_subfamily_4_group_A_member_1__Homo_sapiens_
P12813.1_Nuclear_receptor_subfamily_4_group_A_member_1__Mus_musculus_
XP_022245984.1_probable_nuclear_hormone_receptor_HR38_isoform_X4__Limulus_polyphemus_
CAA75690.1_hormone_receptor_38__Drosophila_melanogaster_
XP_022322351.1_nuclear_receptor_subfamily_4_group_A_member_2_like__partial__Crassostrea_...
isotig15239___ORF_30__frame_3__translation
CAE1263088.1_NR4A2__Sepia_pharaonis_
XP_014786067.1_PREDICTED__probable_nuclear_hormone_receptor_HR38__Octopus_bimaculoides_
isotig22687___BLAST_Hit_translation
Group 4
XP_022305996.1_probable_nuclear_hormone_receptor_HR3_isoform_X2__Crassostrea_virginica_
XP_029653424.1_probable_nuclear_hormone_receptor_HR3__Octopus_sinensis_
NP_730322.1_Ecdysone_induced_protein_75B__isoform_C__Drosophila_melanogaster_
XP_022247851.1_ecdysone_induced_protein_78C_like_isoform_X1__Limulus_polyphemus_
XP_022320890.1_ecdysone_induced_protein_78C_like__Crassostrea_virginica_
XP_012935301.1_ecdysone_induced_protein_78C__Aplysia_californica_
isotig07106___ORF_35__frame_1__translation
XP_029650940.2_nuclear_receptor_subfamily_1_group_D_member_2_like_isoform_X1__Octopus_si...
XP_014779933.1_nuclear_receptor_subfamily_1_group_D_member_2_like_isoform_X1__Octopus_bi...
XP_022324609.1_thyroid_hormone_receptor_beta_like__Crassostrea_virginica_
XP_013790570.1_ecdysone_receptor_like__partial__Limulus_polyphemus_
AAL13804.1_LD26915p__Drosophila_melanogaster_
NP_724459.1_ecdysone_receptor__isoform_C__Drosophila_melanogaster_
XP_014782877.1_PREDICTED__ecdysone_receptor_like__Octopus_bimaculoides_
XP_029636306.1_ecdysone_receptor__Octopus_sinensis_
isotig27046___ORF_1__frame_2__translation
CAE1320634.1_NR1H1__Sepia_pharaonis_
XP_035824311.1_retinoic_acid_receptor_gamma__Aplysia_californica_
XP_022324317.1_retinoic_acid_receptor_alpha_A_like_isoform_X1__Crassostrea_virginica_
XP_014781601.1_PREDICTED__retinoic_acid_receptor_gamma_A_like__partial__Octopus_bimaculo...
XP_029639781.1_retinoic_acid_receptor_gamma_like__Octopus_sinensis_
A2T928.1_Retinoic_acid_receptor_gamma_B__Danio_rerio_
Q91392.2_Retinoic_acid_receptor_gamma_A__Danio_rerio_
P28699.2_Retinoic_acid_receptor_gamma__Xenopus_laevis_
P13631.1_Retinoic_acid_receptor_gamma__Homo_sapiens_
P18911.3_Retinoic_acid_receptor_gamma__Mus_musculus_
P10826.2_Retinoic_acid_receptor_beta__Homo_sapiens_
P22605.1_Retinoic_acid_receptor_beta__Mus_musculus_
Q90271.2_Retinoic_acid_receptor_alpha_A__Danio_rerio_
Q7ZTI3.1_Retinoic_acid_receptor_alpha_B__Danio_rerio_
P51126.1_Retinoic_acid_receptor_alpha__Xenopus_laevis_
P11416.1_Retinoic_acid_receptor_alpha__Mus_musculus_
P10276.2_Retinoic_acid_receptor_alpha__Homo_sapiens_
Group 1
2.0

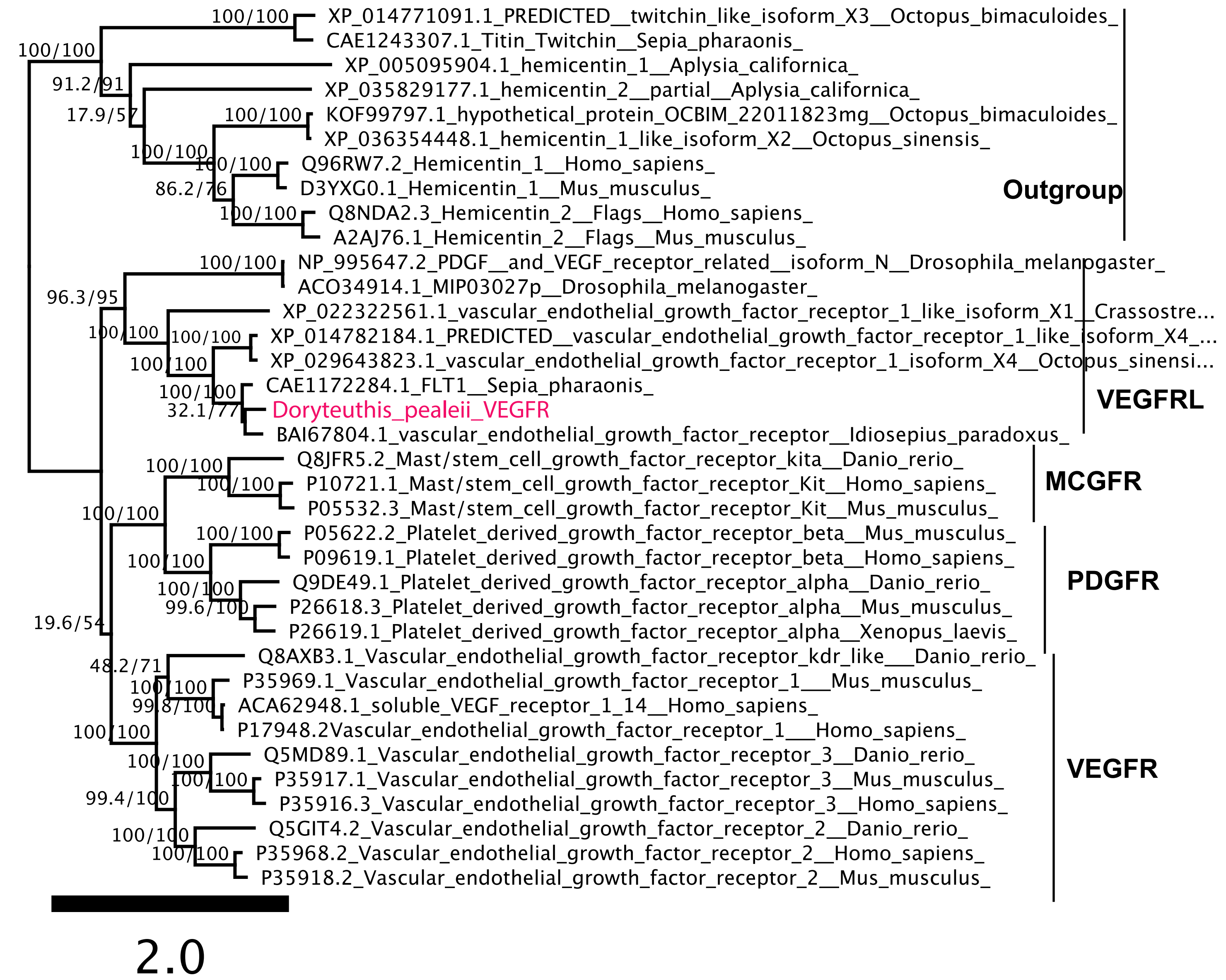
XP_014771091.1_PREDICTED__twitchin_like_isoform_X3__Octopus_bimaculoides_
CAE1243307.1_Titin_Twitchin__Sepia_pharaonis_
XP_005095904.1_hemicentin_1__Aplysia_californica_
XP_035829177.1_hemicentin_2__partial__Aplysia_californica_
KOF99797.1_hypothetical_protein_OCBIM_22011823mg__Octopus_bimaculoides_
XP_036354448.1_hemicentin_1_like_isoform_X2__Octopus_sinensis_
Q96RW7.2_Hemicentin_1__Homo_sapiens_
D3YXG0.1_Hemicentin_1__Mus_musculus_
Q8NDA2.3_Hemicentin_2__Flags__Homo_sapiens_
A2AJ76.1_Hemicentin_2__Flags__Mus_musculus_
NP_995647.2_PDGF__and_VEGF_receptor_related__isoform_N__Drosophila_melanogaster_
ACO34914.1_MIP03027p__Drosophila_melanogaster_
XP_022322561.1_vascular_endothelial_growth_factor_receptor_1_like_isoform_X1__Crassostre...
XP_014782184.1_PREDICTED__vascular_endothelial_growth_factor_receptor_1_like_isoform_X4_...
XP_029643823.1_vascular_endothelial_growth_factor_receptor_1_isoform_X4__Octopus_sinensi...
CAE1172284.1_FLT1__Sepia_pharaonis_
Doryteuthis_pealeii_VEGFR
BAI67804.1_vascular_endothelial_growth_factor_receptor__Idiosepius_paradoxus_
Q8JFR5.2_Mast/stem_cell_growth_factor_receptor_kita__Danio_rerio_
P10721.1_Mast/stem_cell_growth_factor_receptor_Kit__Homo_sapiens_
P05532.3_Mast/stem_cell_growth_factor_receptor_Kit__Mus_musculus_
P05622.2_Platelet_derived_growth_factor_receptor_beta__Mus_musculus_
P09619.1_Platelet_derived_growth_factor_receptor_beta__Homo_sapiens_
Q9DE49.1_Platelet_derived_growth_factor_receptor_alpha__Danio_rerio_
P26618.3_Platelet_derived_growth_factor_receptor_alpha__Mus_musculus_
P26619.1_Platelet_derived_growth_factor_receptor_alpha__Xenopus_laevis_
Q8AXB3.1_Vascular_endothelial_growth_factor_receptor_kdr_like___Danio_rerio_
P35969.1_Vascular_endothelial_growth_factor_receptor_1___Mus_musculus_
ACA62948.1_soluble_VEGF_receptor_1_14__Homo_sapiens_
P17948.2Vascular_endothelial_growth_factor_receptor_1___Homo_sapiens_
Q5MD89.1_Vascular_endothelial_growth_factor_receptor_3__Danio_rerio_
P35917.1_Vascular_endothelial_growth_factor_receptor_3__Mus_musculus_
P35916.3_Vascular_endothelial_growth_factor_receptor_3__Homo_sapiens_
Q5GIT4.2_Vascular_endothelial_growth_factor_receptor_2__Danio_rerio_
P35968.2_Vascular_endothelial_growth_factor_receptor_2__Homo_sapiens_
P35918.2_Vascular_endothelial_growth_factor_receptor_2__Mus_musculus_
Outgroup
VEGFRL
MCGFR
PDGFR
VEGFR
100/100
91.2/91
17.9/57
86.2/76
96.3/95
32.1/77
19.6/54
48.2/71
99.8/100
99.6/100
99.4/100
2.0

### Supplemental Figure 2

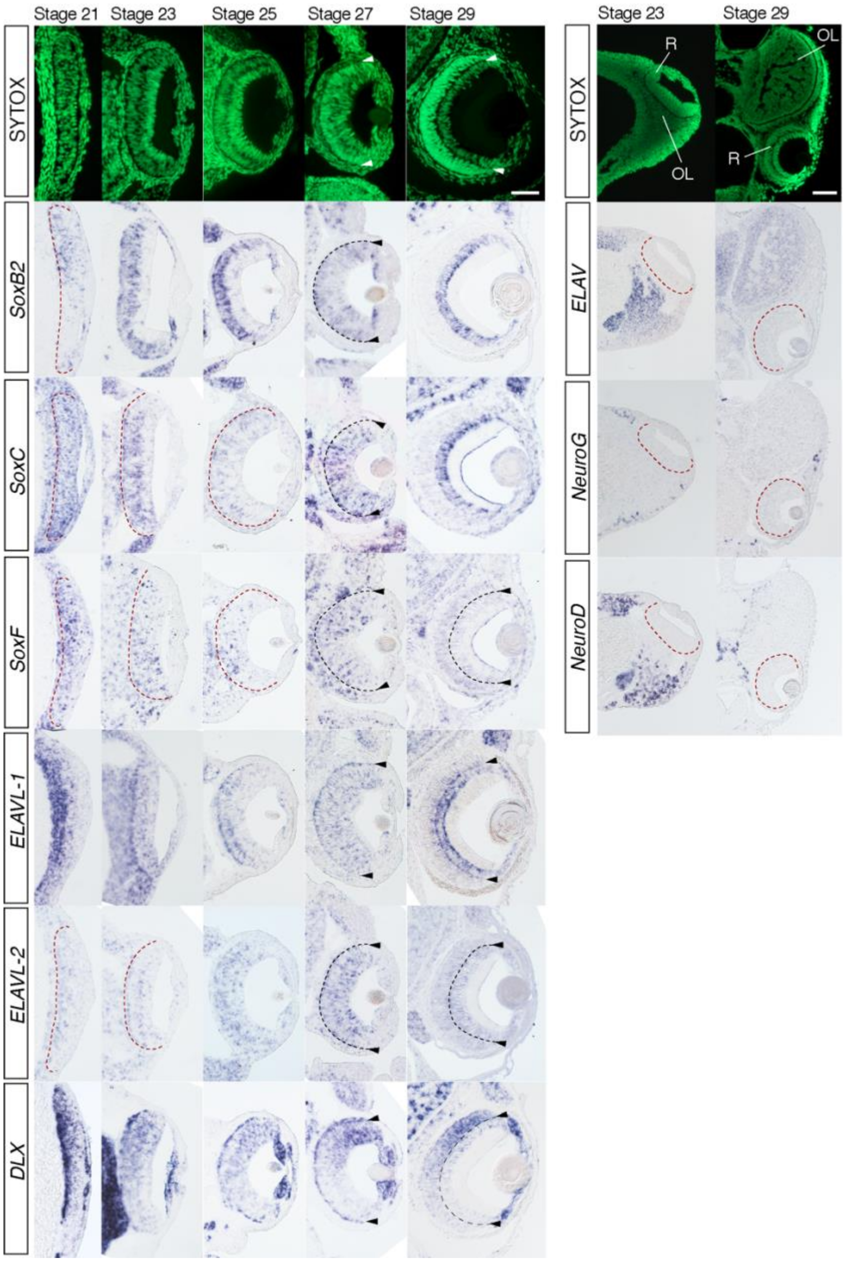


**Supplemental Figure 2: Canonical neurogenesis gene expression in the squid retina throughout development.** *DpSoxB2* stage 23-29 expression recapitulates *DpSoxB1* expression in the retina showing correlation with retinal progenitor cell expression. *DpSoxB2* Stage 21 expression differs from *DpSoxB1* showing a gradient of expression and is not uniform. *DpSoxC* and *DpSoxF* are both expressed in the developing retina throughout development. *DpSoxC* is specifically expressed in the support cell layer at Stage 29. Both *DpELAVL-1* and *DpELAVL-2* is expressed in the developing nervous system and the developing retina. *DpELAV-1* is expressed in the support cell layer and a population of cells posterior of the basal membrane similar to *DpEphR* expression. *DpDLX* expression maintains a gradient throughout development. *DpELAV, DpNeuroG, DpNeuroD* are all expressed in the developing nervous system but are excluded from retinal development. Nuclear SYTOX-Green shown in green. Red dotted lines identify the posterior retinal boundary. Arrowheads and the black and white dotted lines identify the basal membrane. Stage 21, 23, & 25 embryo anterior is down. Stage 27 & 29 embryo dorsal is up. *OL:* Optic Lobe; *R:* Retina. Scale is 50um.

### Supplemental Figure 3

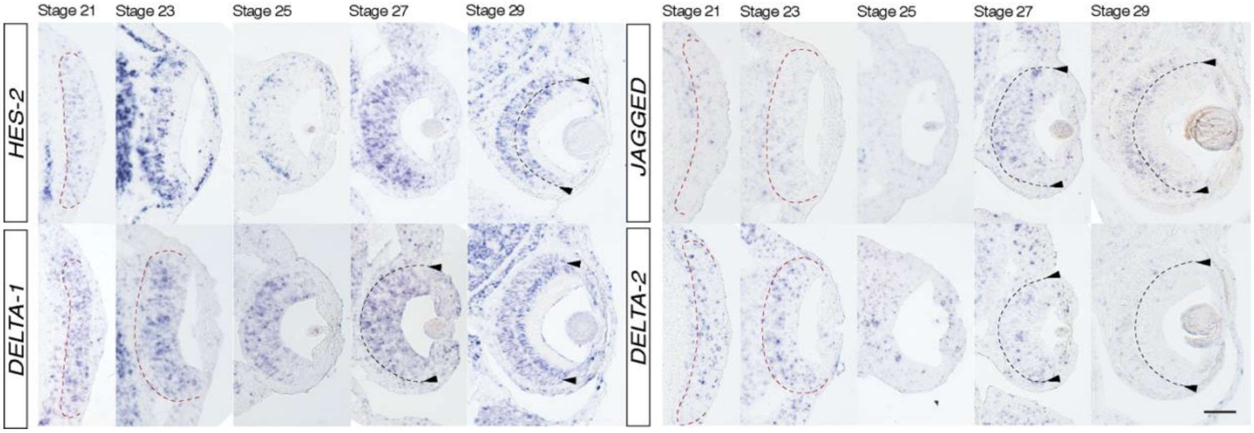


**Supplemental Figure 3: Notch signaling pathways member expression throughout development.** *DpHes-2* shows expression at stage 23 correlated with retinal progenitor cell population. *DpHes-2* expression is enriched in the posterior at stage 25. *DpJagged* expression is found in the retina during development, although inconsistent across stages. *DpJagged* is clearly expressed at stage 27 and 29. *DpDelta-1* is uniformly expressed across the retina at all stages. *DpDelta-2* shows uniform expression across the retina from stage 21 to stage 27. *DpDelta-2* is not expressed at stage 29. Red dotted lines identify the posterior retinal boundary. Arrowheads and the black dotted lines identify the basal membrane.Scale 50um

### Supplemental Figure 4

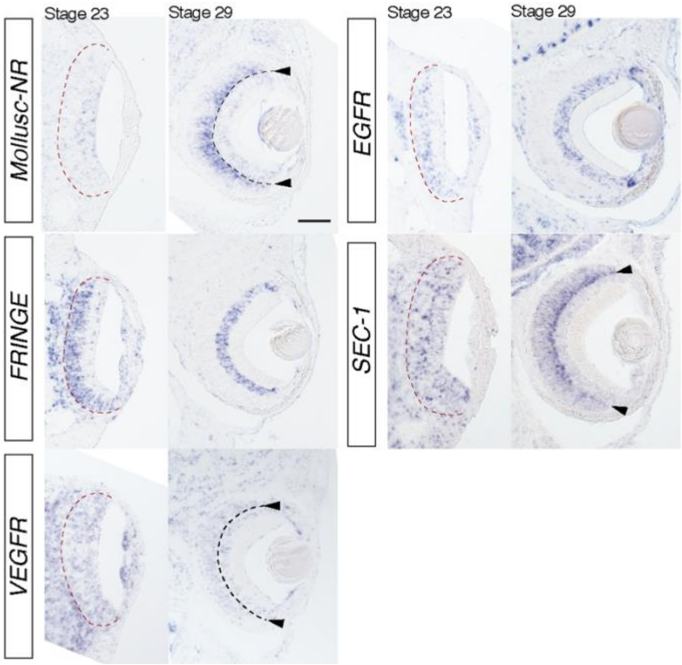


**Supplemental Figure 4: Wildtype expression at stage 23 and 29 of differentially expressed genes identified in DAPT:Control RNA-seq experiment.** To better understand the role of genes that have differential gene expression in our DAPT RNA-seq experiment, we performed wildtype *in situ* hybridization at stage 23 and stage 29, at the start of our DAPT experiment and when cells have differentiated. *DpMollusc-NR* is enriched in the anterior at stage 23 and posterior to the basal membrane at Stage 29. *DpFringe,* a part of the Notch signaling pathway, shows similar expression at stage 23 and 29 as *DpSoxB1, DpHes-1* and *DpNotch,* correlated with the retinal progenitor cell population. *DpVEGFR* is uniformly expressed in the retina at both stage 23 and stage 29. *DpEGFR* expression correlates with *DpSoxB1. DpSec-1* is uniformly expressed at stage 23 and enriched posterior of the basal membrane at Stage 29. Stage 23 embryo anterior is down and at stage 29 embryo dorsal is up. Red dotted lines identify the posterior retinal boundary. Arrowheads and the black and white dotted lines identify the basal membrane. Scale 50um.

### Supplemental Figure 5

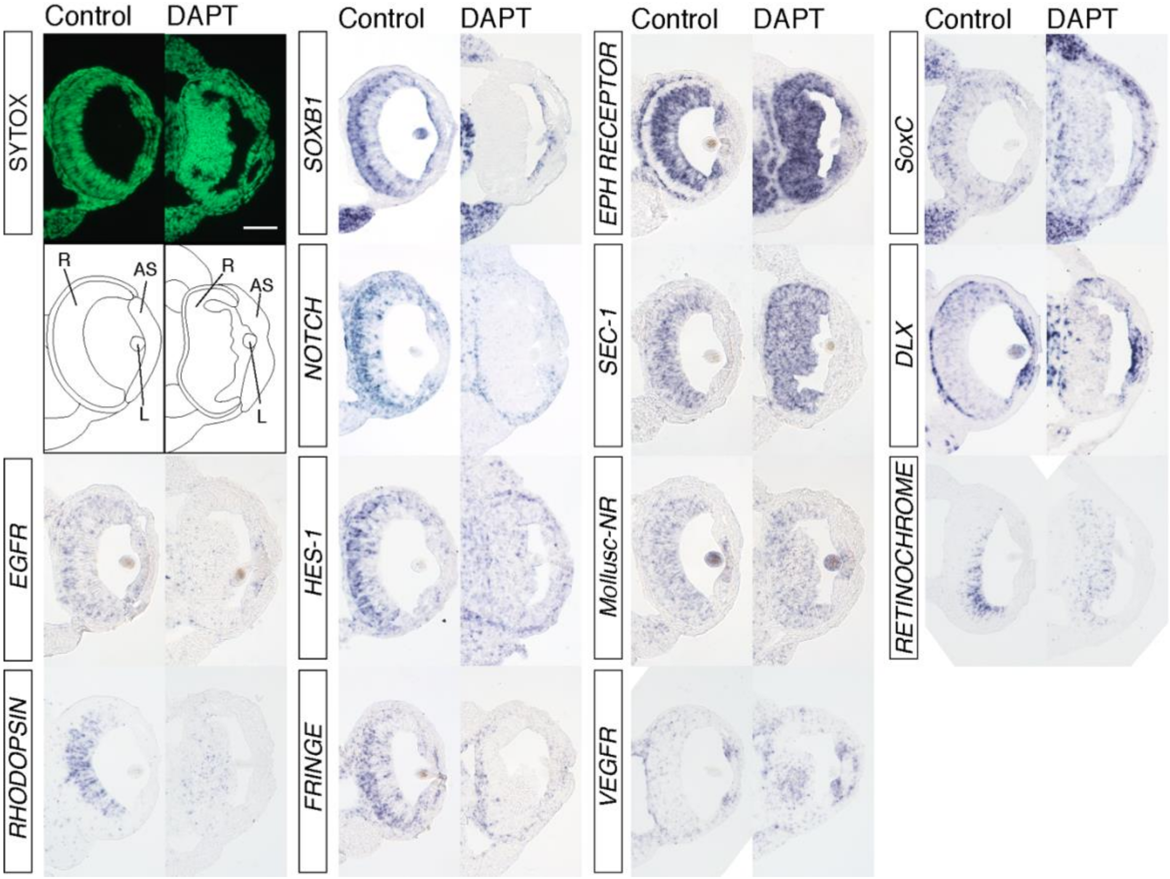


**Supplemental Figure 5: DAPT and Control *in situ* hybridization studies.** Experiments start at stage 23, treated with 20uM DAPT or DMSO for 24 hours. Embryos were fixed, wax embedded and *in situ* hybridization was performed. Nuclear SYTOX-Green shown in green. Disorganization is apparent in DAPT treated retinas. Cartoon with label anatomy below. *R:* Retina; *AS:* Anterior Segment; *L:* Lens. The first and second column show genes with decreased retinal expression. The third column are genes that show increased expression in the retina. The fourth column show changes expression outside the retina (*DpSoxC*) or genes that show similar expression (*DpDlx* and *DpRetinochrome*) after treatment. Scale 50uM.

**Supplemental Table 1: Notch versus control Deseq2 results**

**Supplemental Table 2: Primer Sequences**

| Gene Name | Forward Primer | Reverse Primer |
|---|---|---|
| Delta-1 | AGGGTTTGGTGAACAGTCATCG | TGTCATTTAGGCTGGAAGGGC |
| Delta-2 | CCCTTGGCAGTGTATCTGTGAAG | ACCAAGTTCCCCTGTTACGCAG |
| Jagged | TTTCTTTCCAGCAGTCACCA | CGGTTGGTTAGGAGTCTGAG |
| EGFR | CCATGAGAGTTGCTACCACA | GTATTTCGACCGCCAATGAC |
| EphR | TTTGATAGCCACGGTCATGR | GACAATCCACGAACTCCTCA |
| ELAV | CGGAAACAGCAATGAACACA | ACTCGTGTGCCAAGTAAGAA |
| ELAVL-1 | TAACGGAACCTATTGCCTGG | TGGAGATTCTTGAGACTGCG |
| ELAVL-2 | ATAACGGTAGCCATTCAGCC | RCAAGACAGAGCAACAGGTT |
| HES-1 | GGAAAAACGCAGACGAACACG | GATGCTGTAAACGAAATCACCAGG |
| HES-2 | CGTCTTCGTCAAAAACACAGGTC | GACAACCACCAAAACAGCATCTG |
| NeuroG | CGCTAGTTGTTTCGCCTTAC | CCCACTCCACTGTATCCATC |
| NeuroD | ACGTTGCTCTATCTTGTCCC | ACAGCTTCACCAATCCTCTC |
| Retinochrome | TGTCTGTGGAGTTGTGGTTT | TGCGTGTGGATTCTATGGTT |
| Rhodopsin | TCACGAGAAAGAGATGGCAG | GAGGGGGAGAGGAAAAGTTC |
| Sec-1 | AGCTGTGAAGAAAAATGCCG | TACCCATAGCCAGATCCTGT |
| SoxB1 | CACCATCAGTCGTTGTAGCGTG | GTGAAAAGCAGCCCAAAAAGG |
| SoxB2 | AGAGTCCACCGTTAAGTTCG | TCCTCTCCAAAACGTGTACC |
| SoxC | TGCTTTTTGGTTCGCAACTCG | GGCAAGGTGATTTGATGAGGG |
| SoxF | TGGTGGGTCGGTCAACAGAATAG | CGGGCAATAGAATCATCGCAG |
| Mollusc-NR | TGACTCATCAAACGTGGGGT | ACTGAGTGTACGAAGGGCAG |
| EGFR | CCATGAGAGTTGCTACCACA | GTATTTCGACCGCCAATGAC |
| Fringe | GTCATTGAGCGTTGTAGGCC | AGACAGGGGTTTTATGGGGG |
| VEGFR | CAGTCATTGTTGGCAGTTCC | CATCCGTCGCATATAAGGGT |
